# Reef fishes of praia do Tofo and praia da Barra, Inhambane, Mozambique

**DOI:** 10.1101/139261

**Authors:** Alexander J. Fordyce

## Abstract

The coral reefs around Praia do Tofo and Praia da Barra, southern Mozambique, are known for their aggregations of marine megafauna but few studies have examined their reef fish biodiversity. This study assesses for the first time the ichthyofaunal diversity of the seas around Praia do Tofo and Praia da Barra. Methods involved underwater observations during recreational dives between February and September 2016, and the use of photographic records from 2015. A total of 353 species, representing 79 families, were recorded from 16 patch reefs in the region. The area shows comparable species diversity to others in the southwestern Indian Ocean, suggesting these reefs are in good condition. But high primary productivity driven by coastal upwelling may make fish diversity and trophic structure unreliable measures of the health of these reefs. Future studies investigating the sustainability of this ecosystem would benefit from utilising a wide range of reef health measures.

## Introduction

The ecotourism industry of the Inhambane province in southern Mozambique accounts for approximately 7% of the province’s annual income (Mutimucuio & Meyer, 2011). The primary tourism hotspots are the Bazaruto Archipelago National Park (BANP) and the southern area around the Inhambane peninsula. In the latter, the seas around Praia do Tofo & Praia da Barra (hereafter referred to as PTPB) are particularly important due to their resident populations of manta rays and whale sharks (Pierce *et al*. 2010; Tibirica *et al*. 2011). Venables *et al*. (2016) estimate that manta ray tourism alone contributes $34 million USD per annum to the province’s economy. Scientific research in the PTPB area has thus predominantly focused on these charismatic species (e.g. Rohner *et al*. 2013; 2014); so far, very little research has been conducted on the biodiversity of resident fish populations. This aspect of the PTPB’s marine ecosystem is expected to gain value in the future, as has occurred in the BANP (Schleyer & Celliers, 2005), due to the continued decline of local megafauna populations (Rohner *et al*. 2013). As of 2014, the United Nations & World Heritage Convention (2014) recommend that the protected area currently represented by the BANP be extended south to include the seas around PTPB. Knowledge of the fish biodiversity of this area will help support this recommendation.

Species richness information is currently missing from the PTPB seas but this data is vital for future ecosystem management. Biodiversity data is necessary to identify key biological components (as per Pereira, 2000), provide a baseline from which ecosystem stability and function can be assessed (as per Cleland, 2011), and to predict the effects of biodiversity loss on ecosystem provision (as per Bellwood & Hughes, 2001; Gillibrand, Harries & Mara, 2007; Maggs *et al*., 2010). The PTPB area is bordered by the tropical and sub-tropical latitudes of the southwestern Indian Ocean and are home to a number of different reef habitats likely to support diverse reef fish assemblages. The most common habitats are deepwater, offshore patch reefs which are characteristic of southern Mozambique and typically have low levels of coral cover (e.g. Pereira, 2000; Motta *et al*., 2002; Schleyer & Celliers, 2005). Other marine ecosystems in the region include mangrove swamps, estuarine reefs and shallow inshore fringing reefs. This range of reef and coastal environments provide substantial habitat and nursery grounds for fish species in the area. The PTPB area has a relatively large associated human population of over 250,000 people (Instituto Nacional de Estatística, 2007), based primarily in the cities of Maxixe & Inhambane (Fig. 1). But there is little to no management in place to safeguard the marine ecosystems and the services they provide. This study constitutes a baseline assessment of fish diversity of the reefs surrounding Praia do Tofo & Praia da Barra, and highlights the need for further investigations into the state of these ecosystems.

**Figure 1.**
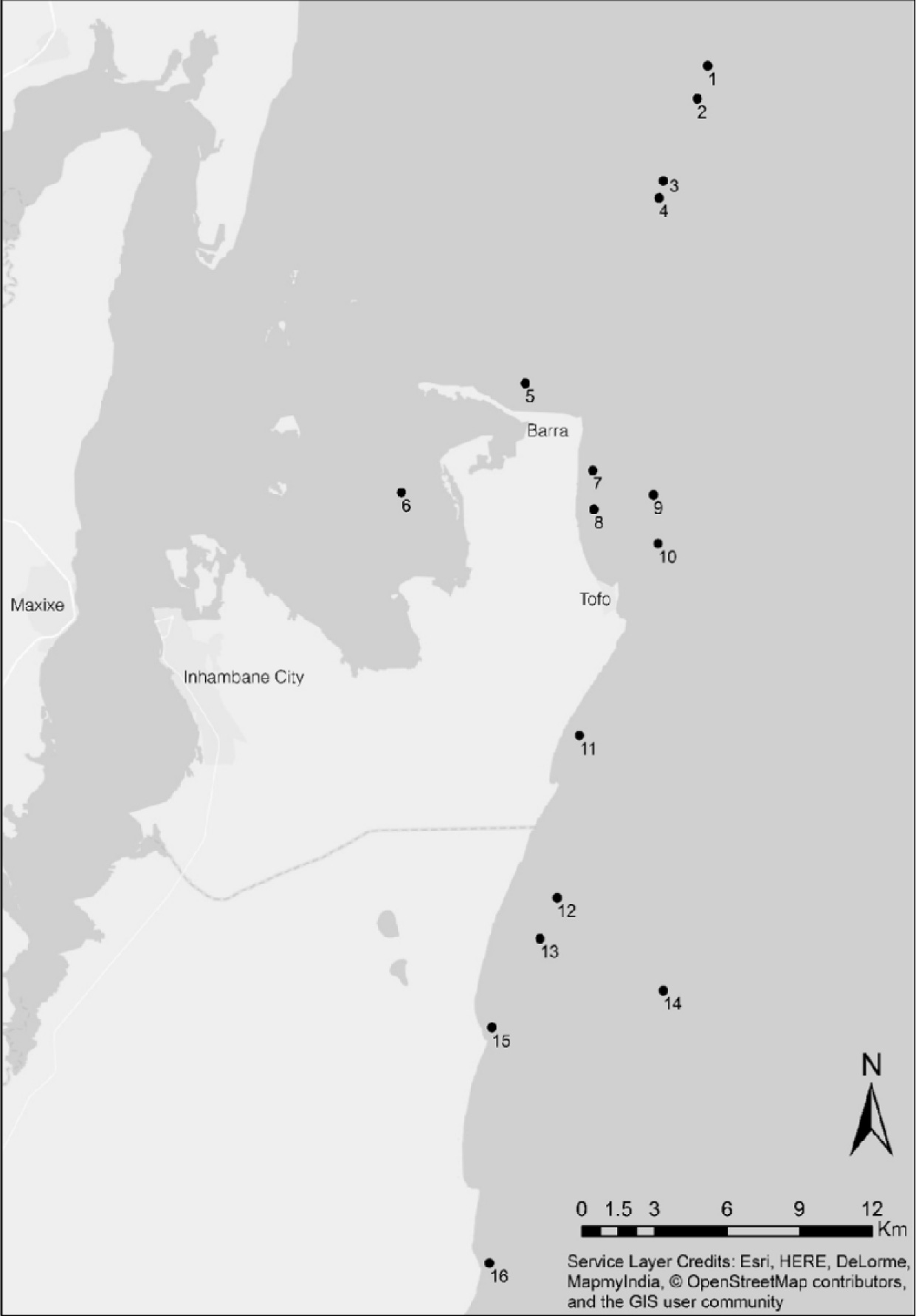
Map of the study area and its location along the coast of Mozambique (inset). Sampled reefs are indicated by (•); their broad characteristics are described in Table 1.

## Materials & Methods

### Study Site

Praia do Tofo (23° 51.205’ S 35° 32.882’ E) and Praia da Barra (23° 47.541’ S 35° 31.142’ E) harbour a number of shallow fringing coral reefs. However, many of the sites frequented by the local dive industry are in deeper waters to the north and south. In this study, diversity was recorded on reefs spanning approximately 40 km along the coast of the Inhambane province (Fig. 1). A total of 16 reef sites between 1 and 32 m (Table 1) were surveyed between February and September 2016.

**Table 1.** Names and descriptions of sampled reefs, including the underwater survey method used and the amount of time spent surveying each location

| Site Name<br>(Number) | Site Description | Sampling<br>Method | Sampling<br>Time<br>(mins) |
| --- | --- | --- | --- |
| Amazon (1) | Offshore, horseshoe reef with an abundance of azooxanthellate soft corals; 23 – 28 metres. | SCUBA | 87 |
| Hospital (2) | Offshore, southward sloping reef with occasional short pinnacles; 24 – 26 metres. | SCUBA | 80 |
| The Office (3) | Topographically complex offshore reef with an abundance of overhangs and valleys with many encrusting soft corals; 22 – 26 metres. | SCUBA | 177 |
| Reggie's (4) | Tall, offshore reef rising between 4 – 8 metres from the seafloor; reef crests are dominated by large colonies of *Tubastrea micranthus*; 22 – 30 metres. | SCUBA | 231 |
| Buddies (5) | Shallow, inshore reef subject to persistent swell and fishing pressure; 8 – 10 metres. | SCUBA | 97 |
| The Wall (6) | Shallow estuarine reef with daily exposure to strong tidal currents; a combination of seagrass, rocky reef and sand patch microhabitats; 0-4 metres. | Snorkel | 70 |
| Mike's<br>Cupboard (7) | Submerged sand dune reef, with many potholes and gullies surrounded by sandy reef flats; 12 – 16 metres. | SCUBA | 108 |
| Salon (8) | Shallow inshore reef composed of multiple large pinnacles surrounded by sandy bottom; subject to high turbidity from wave action; 10-14 metres. | SCUBA | 175 |
| Sherwood<br>Forest (9) | Offshore reef just outside of Tofo bay, made of one large and one smaller pinnacle both supporting large populations of *Tubastrea micranthus*; 22 – 26 metres | SCUBA | 58 |
| Giants Castle<br>(10) | Straight north-south reef with an extensive reef flat and deep reef wall; known within the local dive industry as having the best sighting rate for marine megafauna; 27 – 32 metres. | SCUBA | 214 |
| Marble Arch<br>(11) | Inshore reef exposed to minor wave action; large reef flat with a few large potholes and one large rock arch; 14 – 18 metres. | SCUBA | 51 |
| Rob's Bottom<br>(12) | Very patchy eastward sloping reef that is often subject to high current with high algal cover; 23 – 27 metres. | SCUBA | 158 |
| Manta Reef<br>(13) | A large offshore reef, with a large central reef flat; peripheries are characterised by short, steep reef slopes with a number of tall pinnacles; 18 – 24 metres | SCUBA | 365 |
| Outback (14) | Similar reef shape as Giant's Castle, yet with more small inlets that house a number of deep overhangs and archways; 25 – 30 metres. | SCUBA | 76 |
| Coconut Bay<br>(15) | Shallow inshore rocky reef with small patches of encrusting soft coral and larger swathes of seagrass; 4 – 8 metres. | Snorkel | 53 |
| Paindane<br>Coral Gardens<br>(16) | Small, shallow reef protected from offshore waves by a barrier rock extending from shore; the most abundant coral community in this area, dominated by *Sinularia* spp. soft coral and corymbose acroporids; 1 – 6 metres. | Snorkel | 182 |

### Sampling

The primary method was underwater observations during a random swim. Species were identified *in situ* if possible and recorded on an underwater PVC slate. If required, a photograph was taken for subsequent species identification. Deep sites (> 8 m) were surveyed on SCUBA, as part of a recreational dive charter operated by Peri-Peri Divers. Shallow sites were assessed by snorkelling. Fifty-four individual surveys, totalling 2218 minutes of observation time were undertaken (total surveying times for each site are shown in Table 1). The species richness recorded from underwater observations was supplemented through the inclusion of species that had been sighted in the year preceding the survey period, and for which there was photographic evidence available from local ecotourism and dive operators (e.g. *Mola mola*). The inclusion of solicited data outside the study period was conducted to represent rare or seasonally restricted species. Data collection was approved by the Maritime Administration of the City of Inhambane and the Ministry of Justice.

### Estimated richness and regional comparisons

To determine the number of conspicuous species missed during the visual census, the Coral Fish Diversity Index (CFDI) developed by Allen & Werner (2002) was calculated and compared to the recorded species richness (SR_obs_). The CFDI examines the diversity of six common and easily observable families as representatives of reef fish species richness. These families are Acanthuridae, Chaetodontidae, Labridae, Pomacanthidae, Pomacentridae & Scaridae. In areas < 2000 km^2^, a theoretical species richness (SR_theor_) is then generated using the equation SR_theor_ = 3.39(CFDI) – 20.595 (Allen & Werner, 2002). SR_theor_ was calculated for other reef systems in the southwestern Indian Ocean, using published literature, to draw loose comparisons between the richness of these areas and that observed in the current study (as per Wickel *et al*. 2014).

## Results

A total of 353 species, representing 79 families, were recorded in the current study from 328 visual observations and 25 past photographic records (Table 2). Of the total number of species recorded, 27 were cartilaginous fish and 326 were bony fish. The CFDI-generated Twelve families represented over half of the total recorded diversity, these included Acanthuridae (17), Balistidae (11), Carangidae (10), Chaetodontidae (18), Holocentridae (10), Labridae (32), Lutjanidae (12), Muraenidae (14), Pomacentridae (21), Scorpaenidae (13), Serranidae (19), and Tetraodontidae (10). Nearly half the recorded families (48%) were represented by one species only. Five of these families are monospecific including, Rachycentridae, Rhincodontidae, Rhinidae, Stegostomatidae, and Zanclidae. The most species-rich genera were *Chaetodon* (12), *Epinephelus* (10) and *Gymnothorax* (10).

**Table 2.**
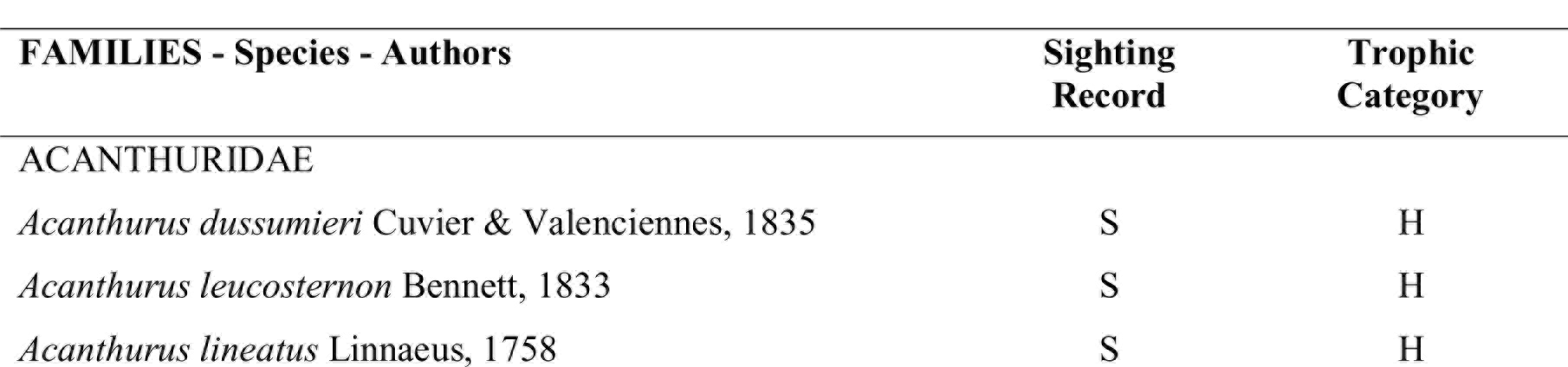
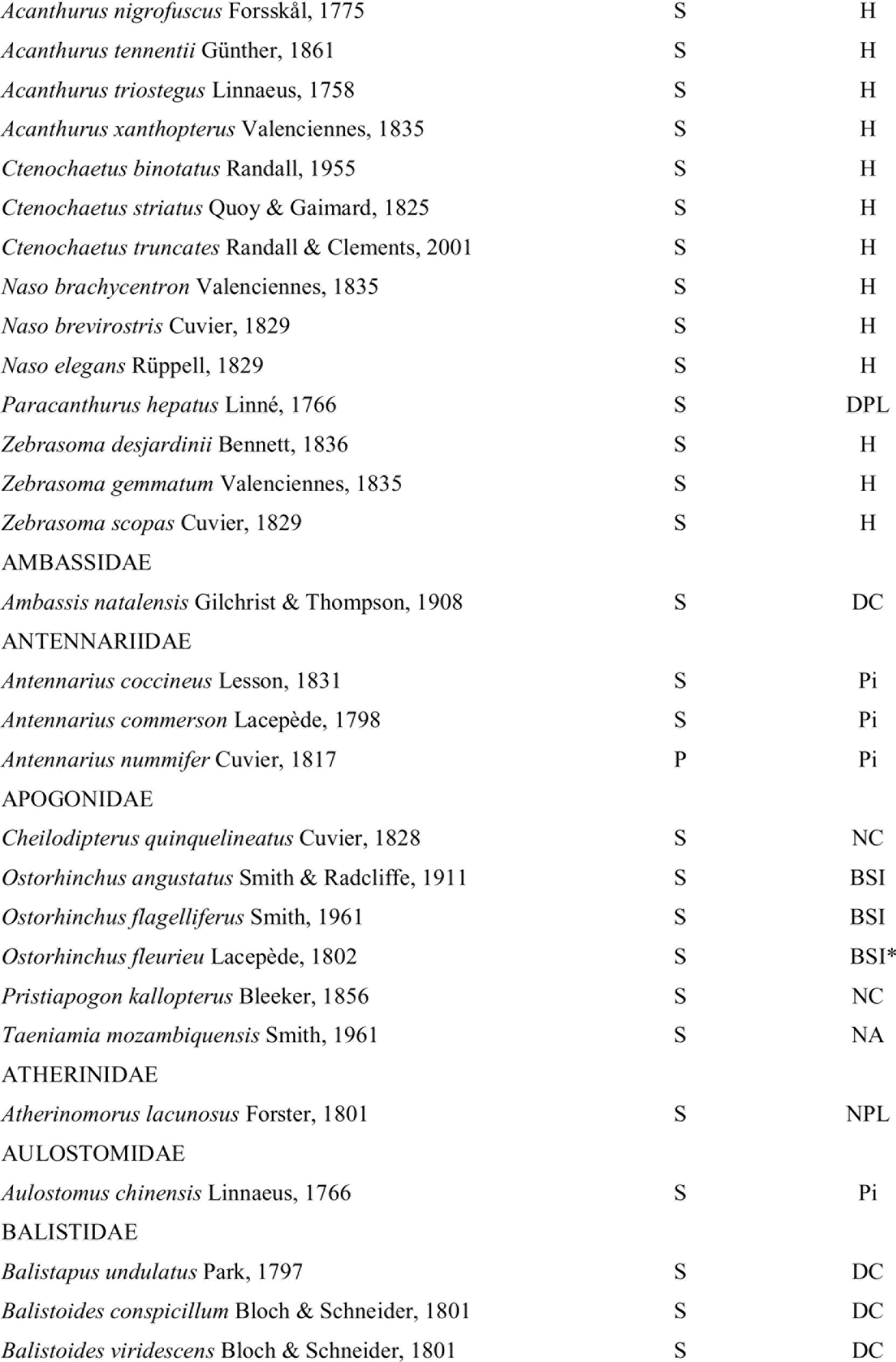
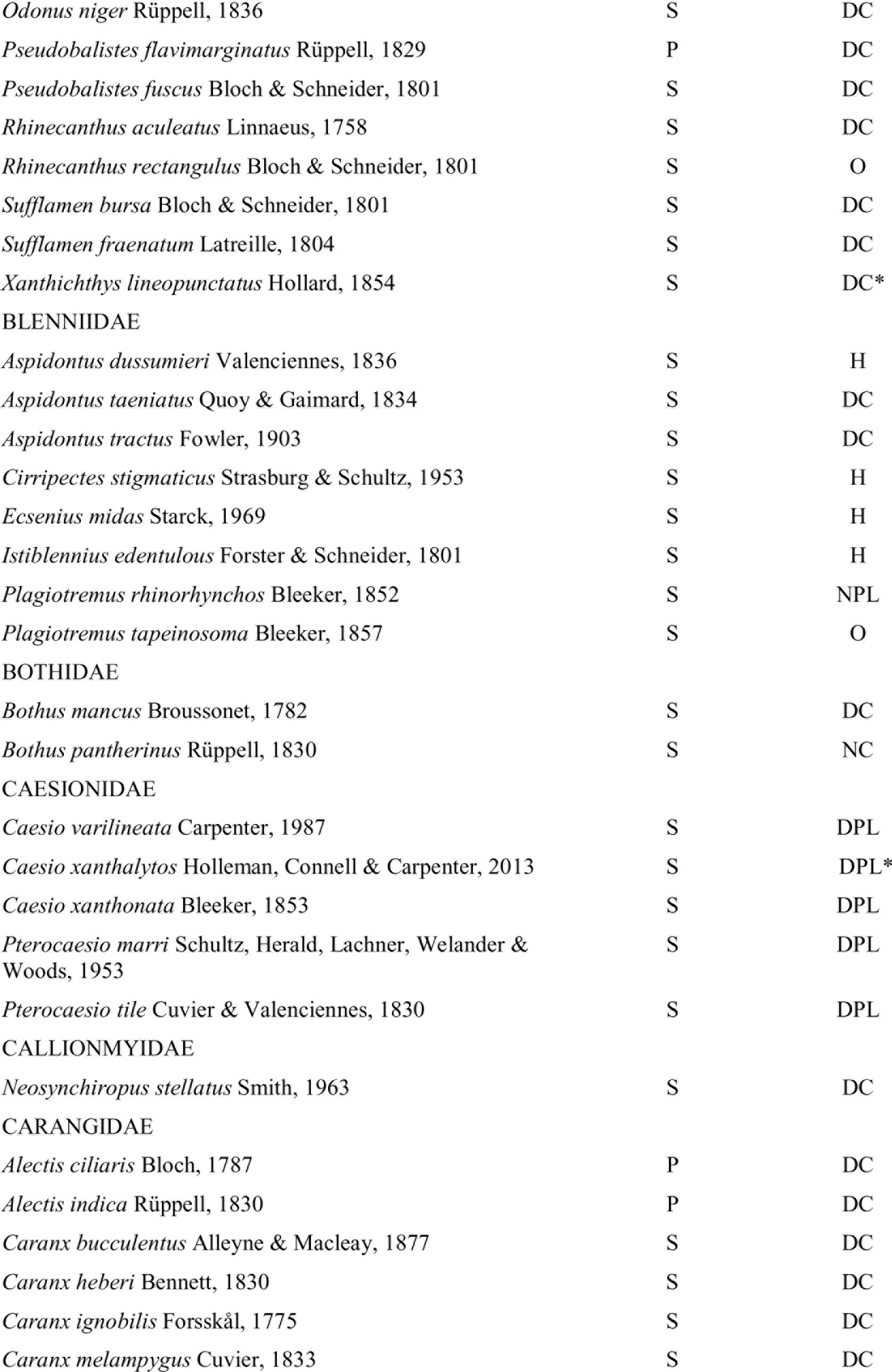
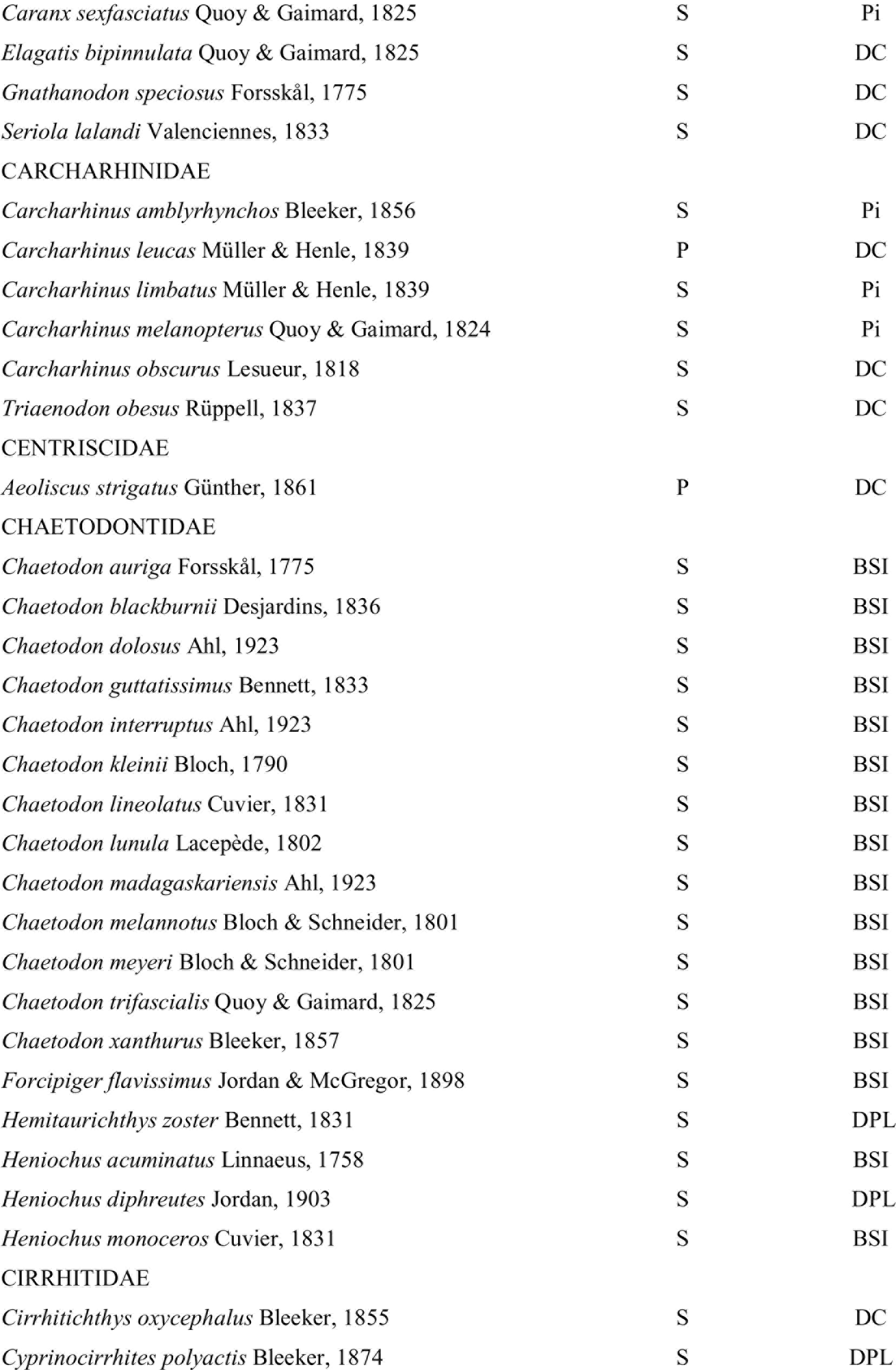
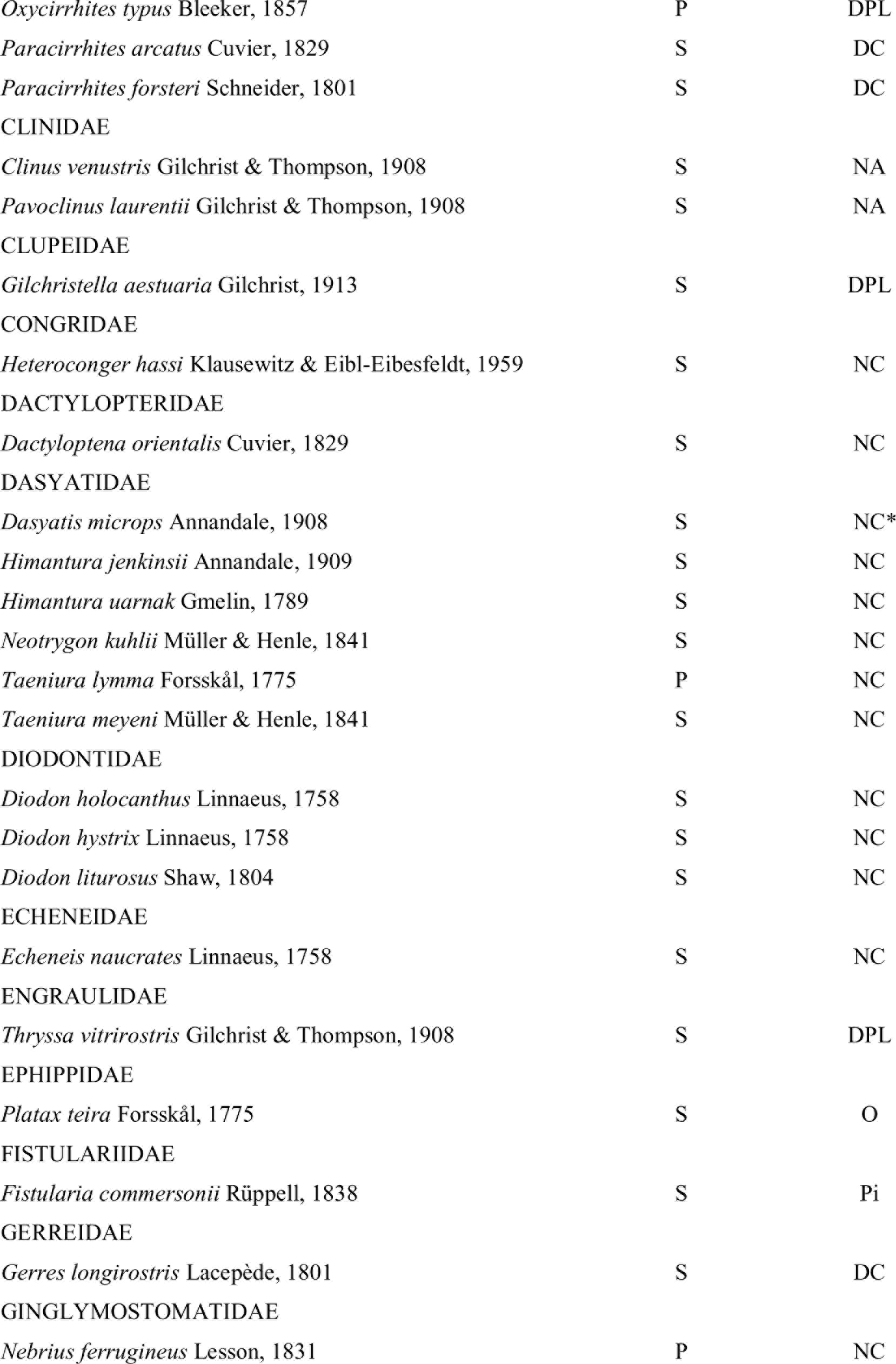
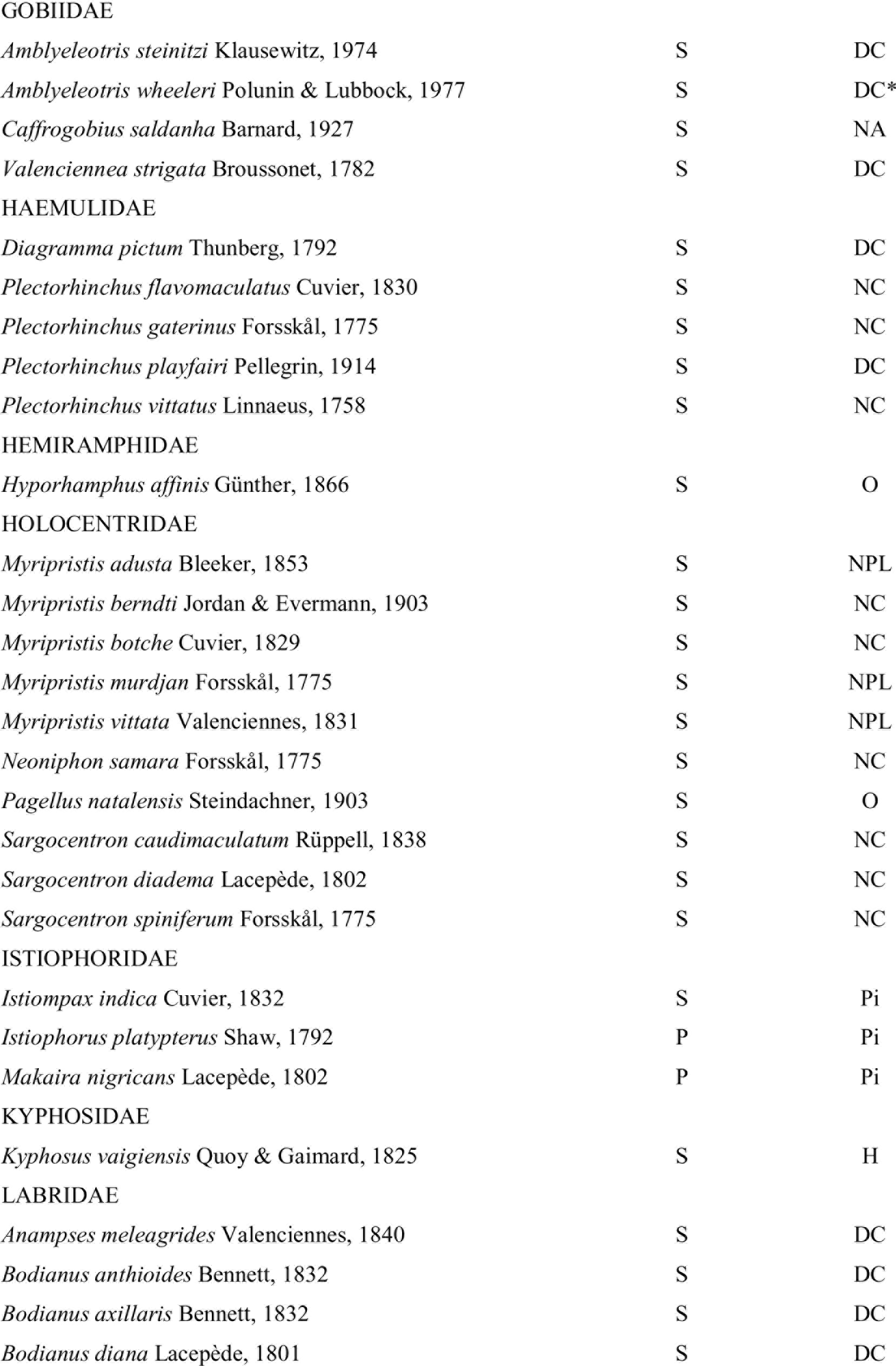
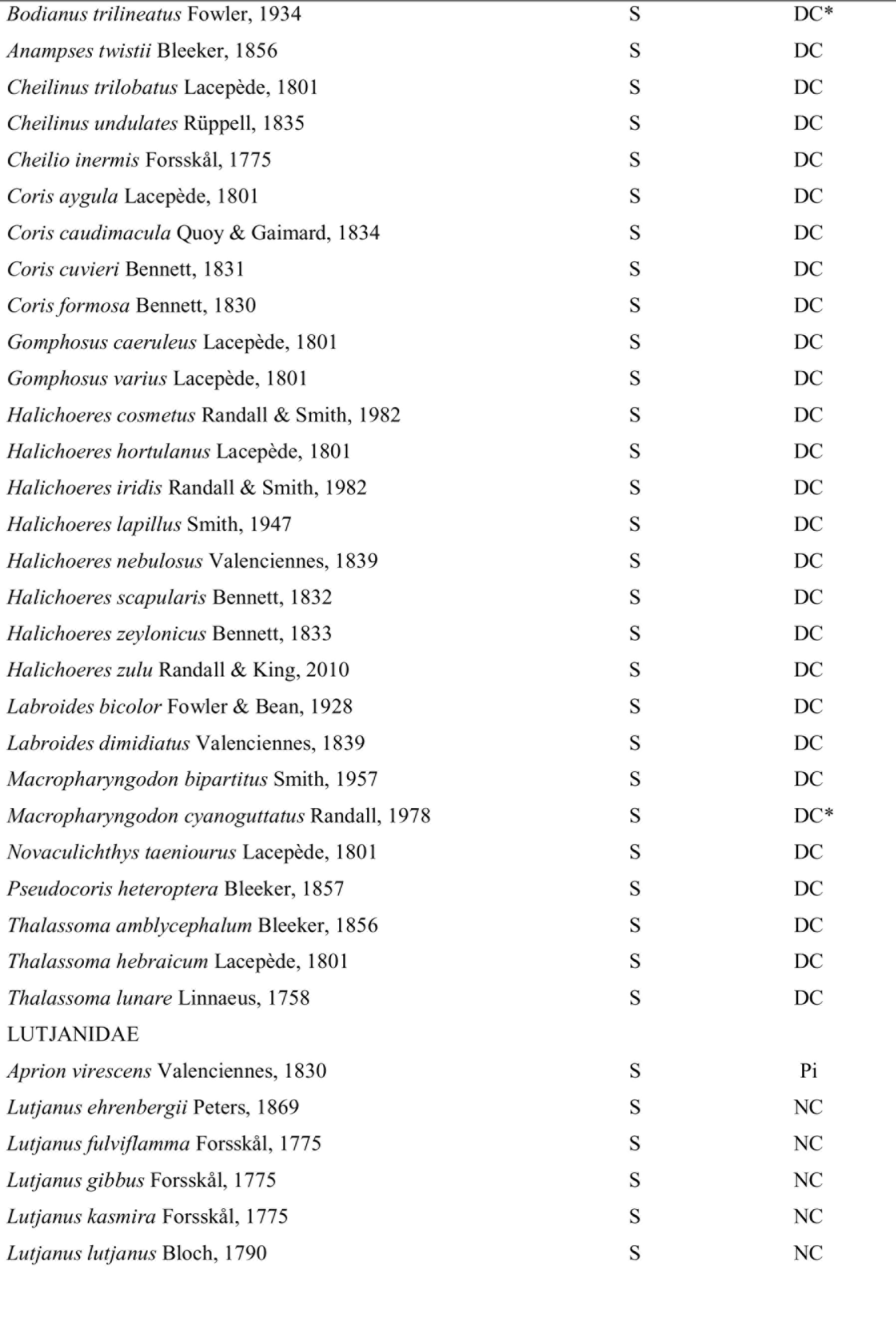
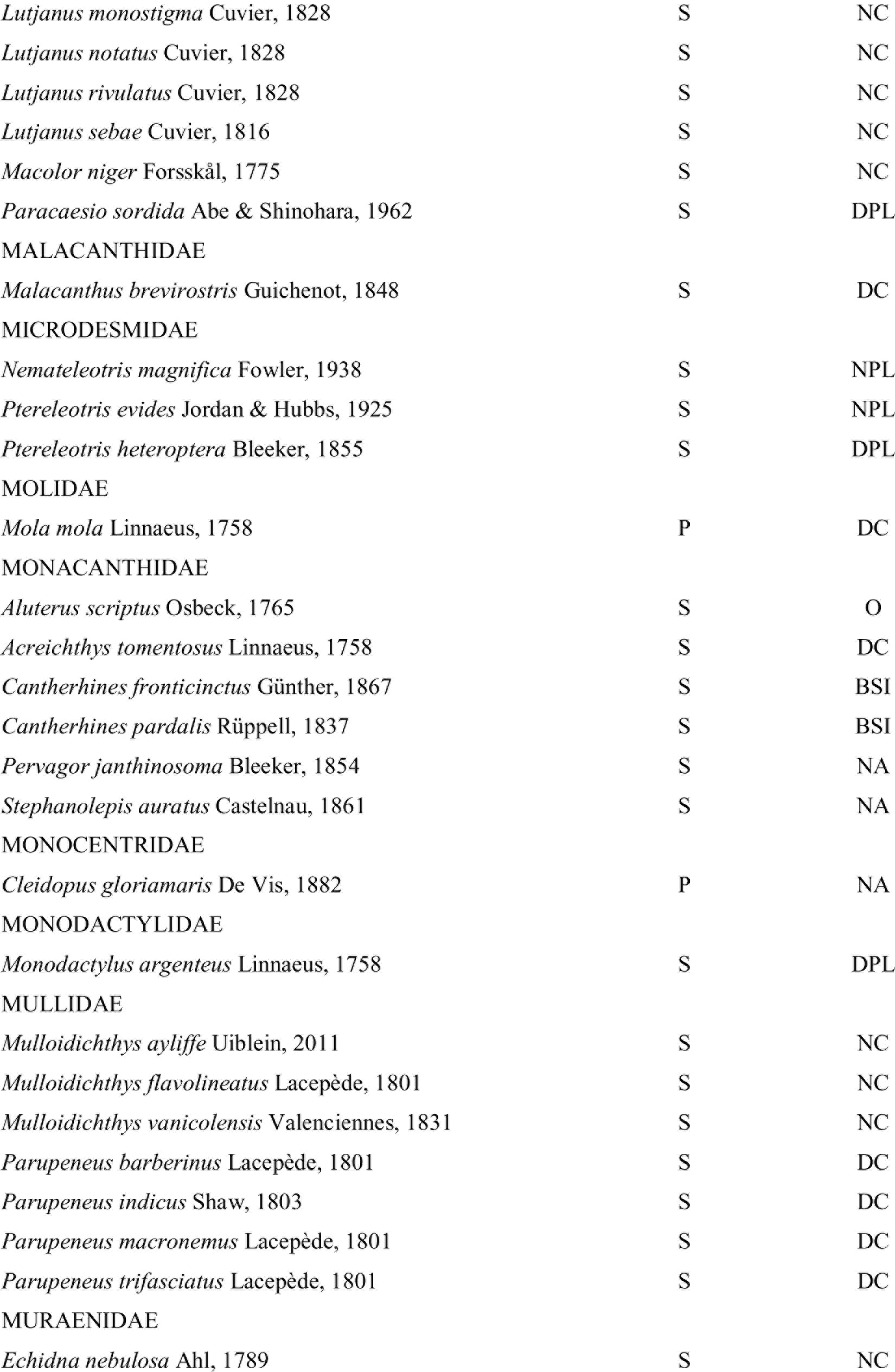
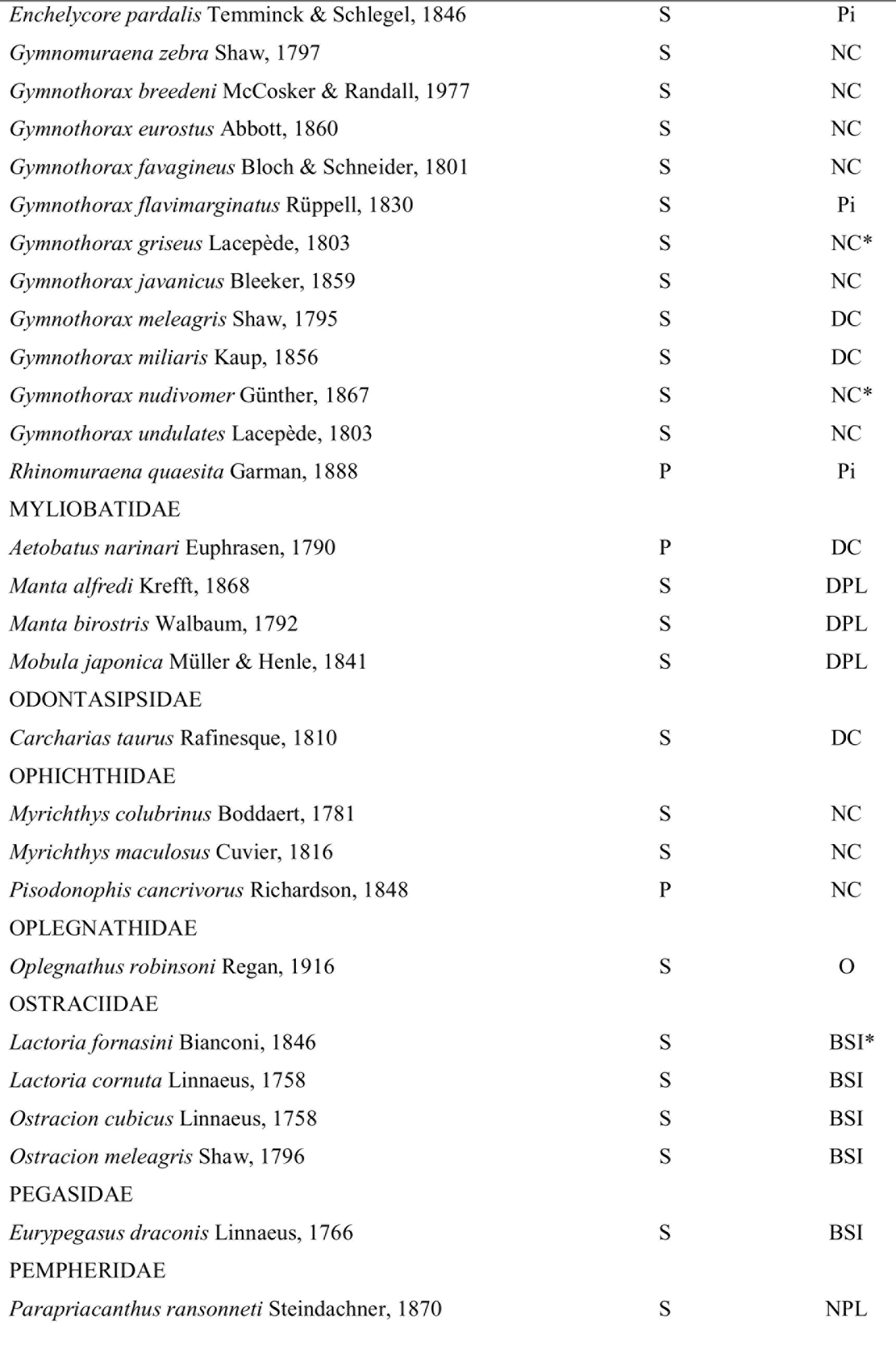
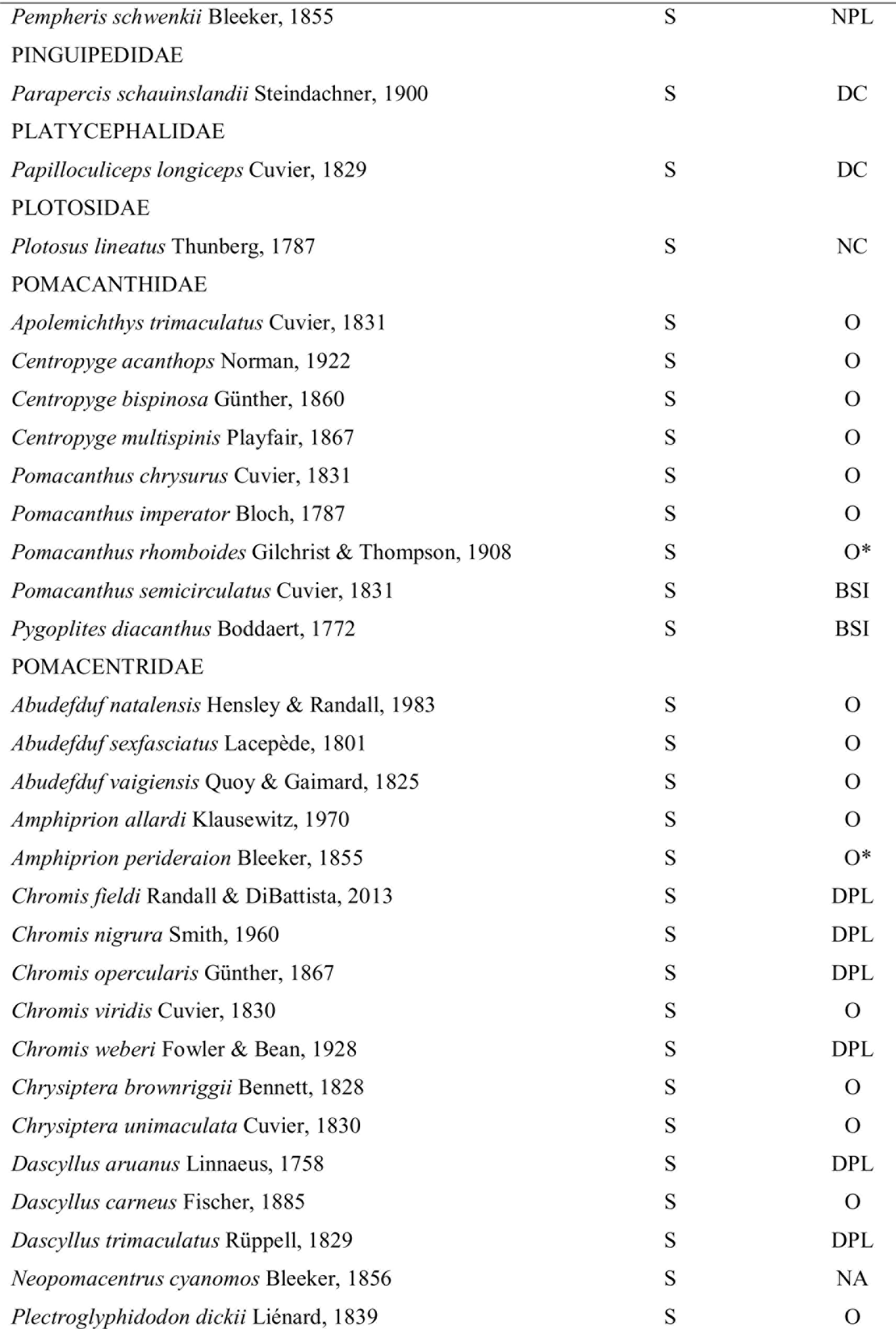
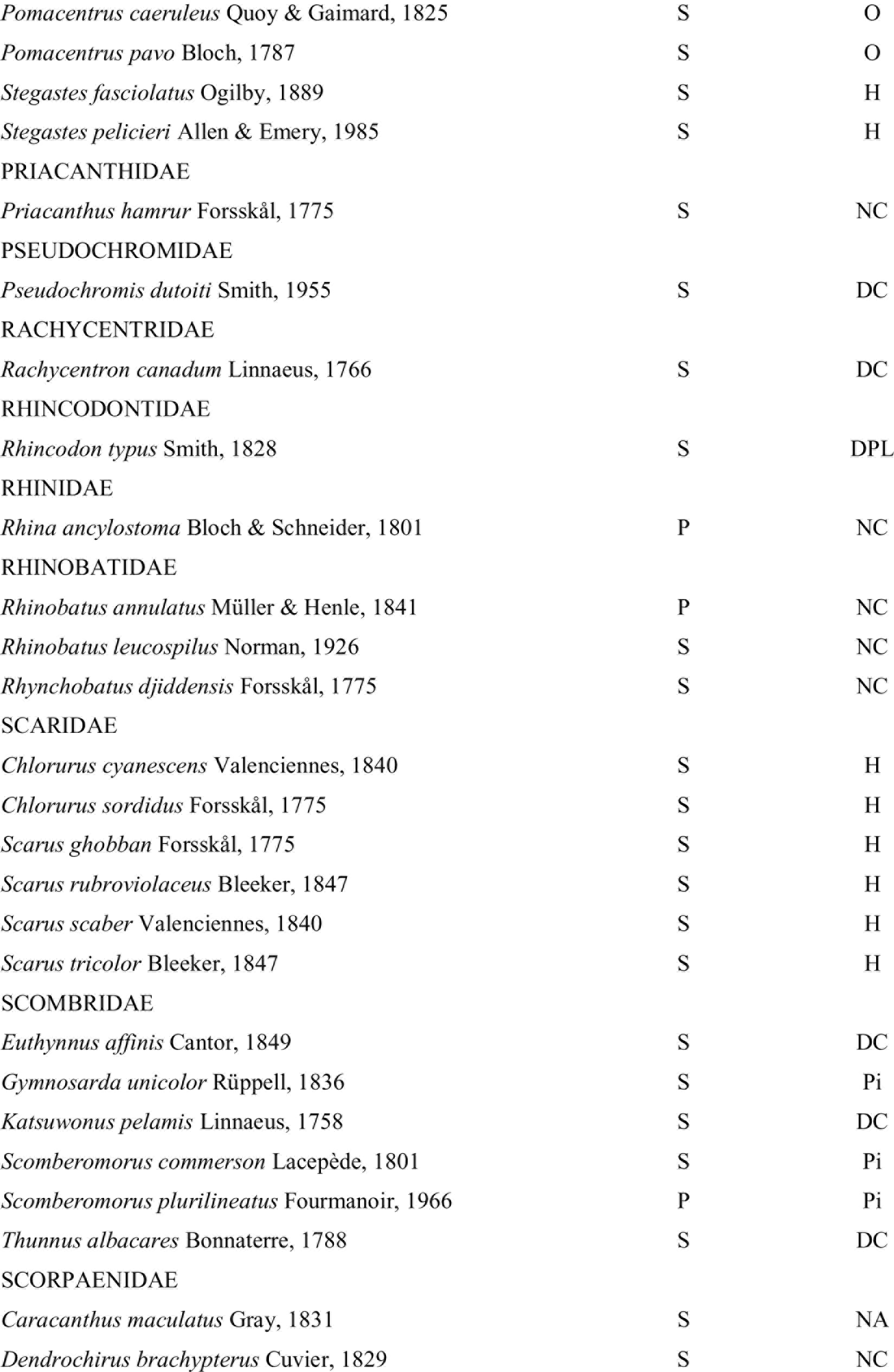
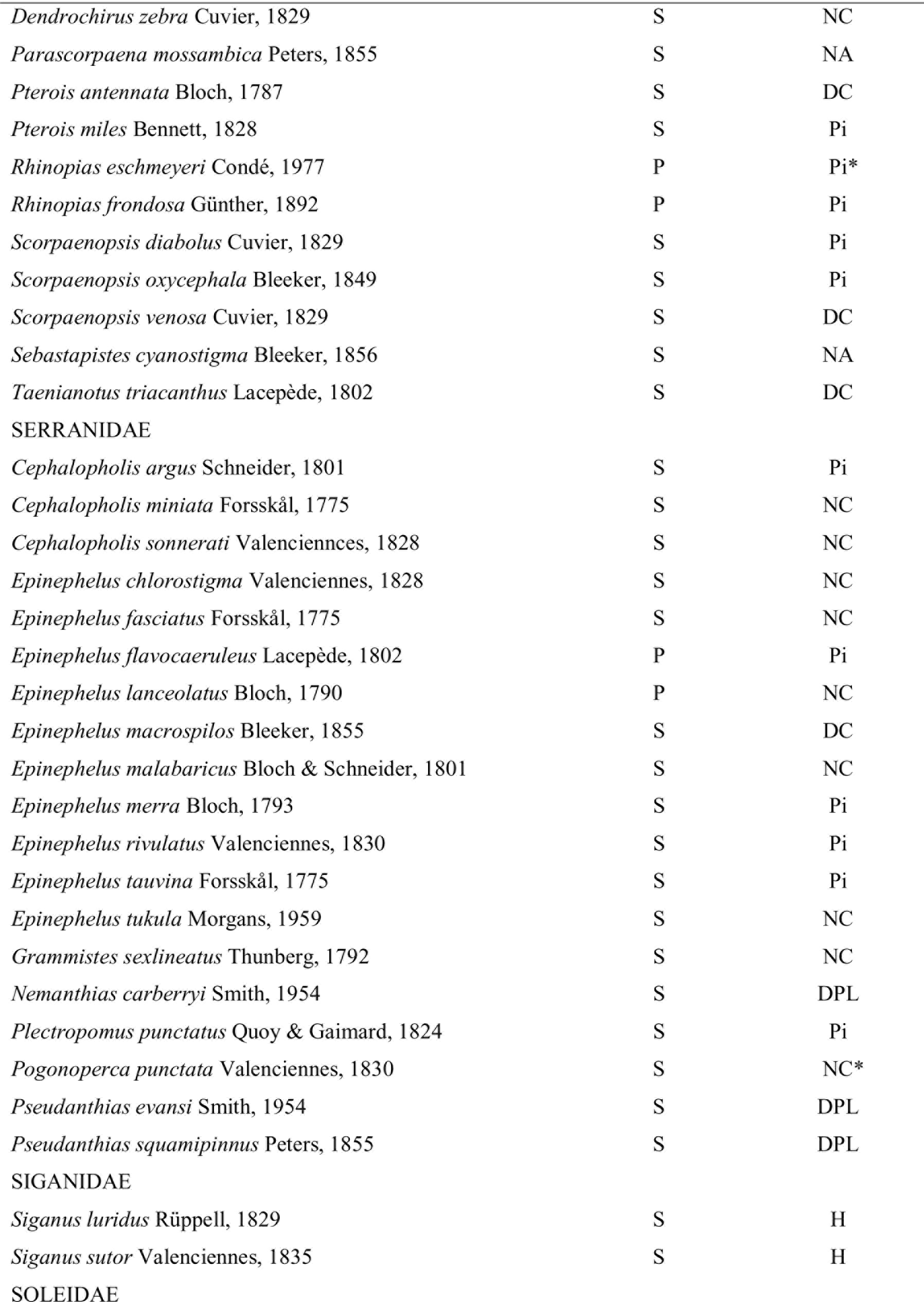
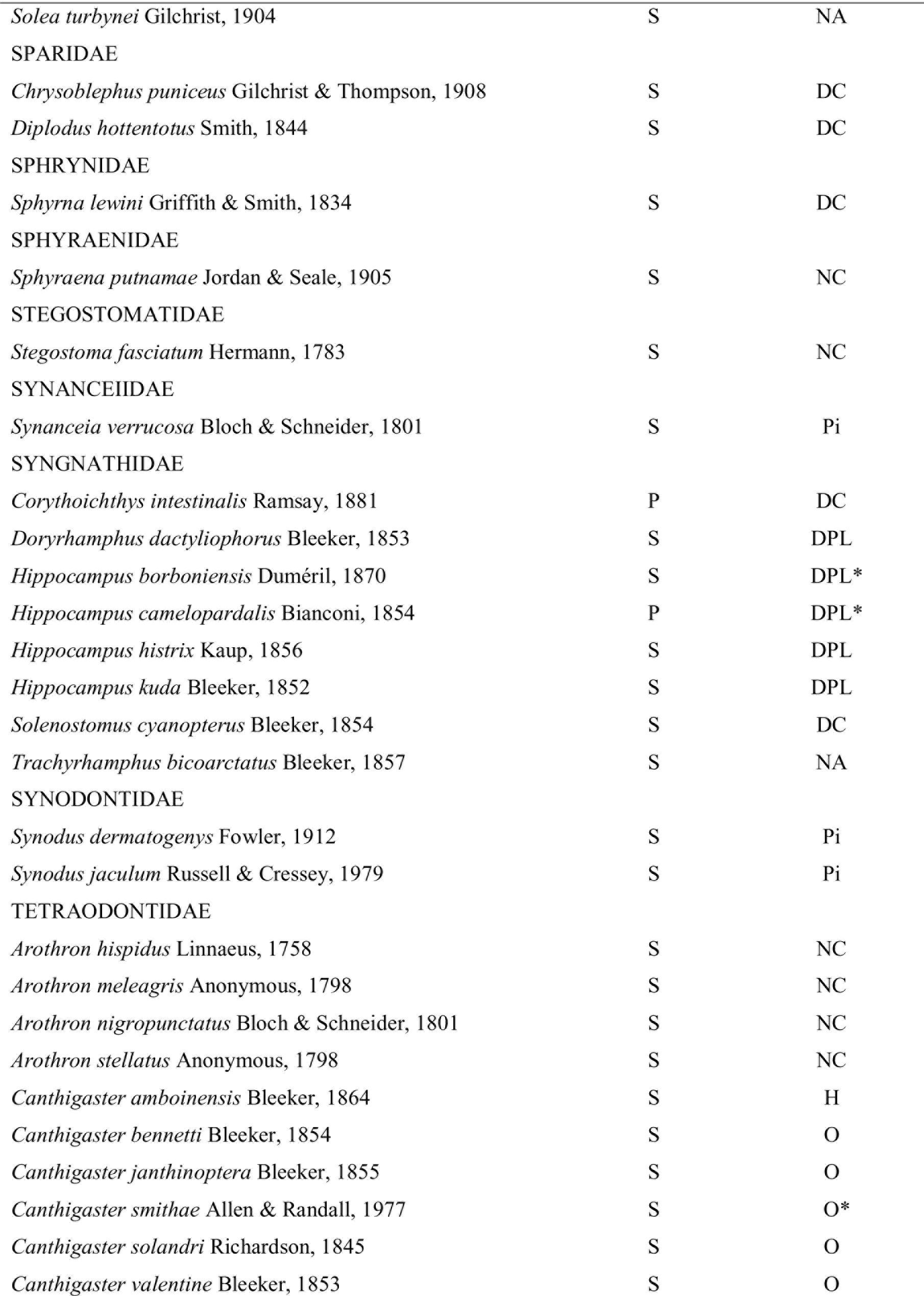
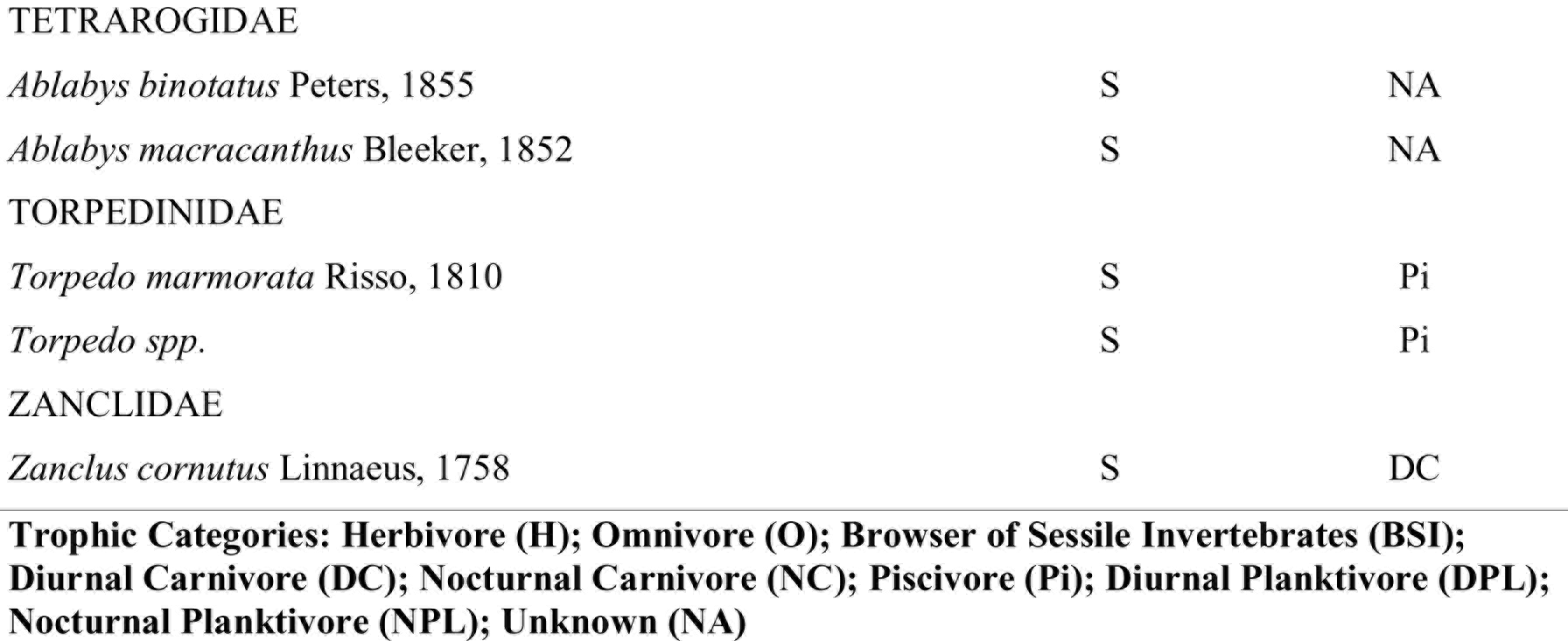
Reef fish species checklist from the PTPB area of Mozambique, sighted through surveys (S) and photographic records (P). Where a species’ trophic category has been assumed from a congener species, it is labelled with a ‘*’. SR_theor_ was 329, lower than the observed species richness (Table 3)

| FAMILIES - Species - Authors | Sighting Record | Trophic Category |
| --- | --- | --- |
| ACANTHURIDAE |  |  |
| *Acanthurus dussumieri* Cuvier & Valenciennes, 1835 | S | H |
| *Acanthurus leucosternon* Bennett, 1833 | S | H |
| *Acanthurus lineatus* Linnaeus, 1758 | S | H |

| **FAMILIES - Species - Authors** | **Sighting Record** | **Trophic Category** |
| --- | --- | --- |
| *Acanthurus nigrofuscus* Forsskål, 1775 | S | H |
| *Acanthurus tennentii* Günther, 1861 | S | H |
| *Acanthurus triostegus* Linnaeus, 1758 | S | H |
| *Acanthurus xanthopterus* Valenciennes, 1835 | S | H |
| *Ctenochaetus binotatus* Randall, 1955 | S | H |
| *Ctenochaetus striatus* Quoy & Gaimard, 1825 | S | H |
| *Ctenochaetus truncates* Randall & Clements, 2001 | S | H |
| *Naso brachycentron* Valenciennes, 1835 | S | H |
| *Naso brevirostris* Cuvier, 1829 | S | H |
| *Naso elegans* Rüppell, 1829 | S | H |
| *Paracanthurus hepatus* Linné, 1766 | S | DPL |
| *Zebrasoma desjardini* Bennett, 1836 | S | H |
| *Zebrasoma gemmatum* Valenciennes, 1835 | S | H |
| *Zebrasoma scopas* Cuvier, 1829 | S | H |
| **AMBASSIDAE** |  |  |
| *Ambassis natalensis* Gilchrist & Thompson, 1908 | S | DC |
| **ANTENNARIIDAE** |  |  |
| *Antennarius coccineus* Lesson, 1831 | S | Pi |
| *Antennarius commerson* Lacepède, 1798 | S | Pi |
| *Antennarius nummifer* Cuvier, 1817 | P | Pi |
| **APOGONIDAE** |  |  |
| *Cheilodipterus quinquelineatus* Cuvier, 1828 | S | NC |
| *Ostorhinchus angustatus* Smith & Radcliffe, 1911 | S | BSI |
| *Ostorhinchus flagelliferus* Smith, 1961 | S | BSI |
| *Ostorhinchus fleurieu* Lacepède, 1802 | S | BSI* |
| *Pristiapogon kallopterus* Bleeker, 1856 | S | NC |
| *Taeniamia mozambiquensis* Smith, 1961 | S | NA |
| **ATHERINIDAE** |  |  |
| *Atherinomorus lacunosus* Forster, 1801 | S | NPL |
| **AULOSTOMIDAE** |  |  |
| *Aulostomus chinensis* Linnaeus, 1766 | S | Pi |
| **BALISTIDAE** |  |  |
| *Balistapus undulatus* Park, 1797 | S | DC |
| *Balistoides conspicillum* Bloch & Schneider, 1801 | S | DC |
| *Balistoides viridescens* Bloch & Schneider, 1801 | S | DC |
| *Odonus niger* Rüppell, 1836 | S | DC |
| *Pseudobalistes flavimarginatus* Rüppell, 1829 | P | DC |
| *Pseudobalistes fuscus* Bloch & Schneider, 1801 | S | DC |
| *Rhinecanthus aculeatus* Linnaeus, 1758 | S | DC |
| *Rhinecanthus rectangulus* Bloch & Schneider, 1801 | S | O |
| *Sufflamen bursa* Bloch & Schneider, 1801 | S | DC |
| *Sufflamen fraenatum* Latreille, 1804 | S | DC |
| *Xanthichthys lineopunctatus* Hollard, 1854 | S | DC* |
| **BLENNIIDAE** |  |  |
| *Aspidontus dussumieri* Valenciennes, 1836 | S | H |
| *Aspidontus taeniatus* Quoy & Gaimard, 1834 | S | DC |
| *Aspidontus tractus* Fowler, 1903 | S | DC |
| *Cirripectes stigmaticus* Strasburg & Schultz, 1953 | S | H |
| *Ecsenius midas* Starck, 1969 | S | H |
| *Istiblennius edentulous* Forster & Schneider, 1801 | S | H |
| *Plagiotremus rhinorhynchus* Bleeker, 1852 | S | NPL |
| *Plagiotremus tapeinosoma* Bleeker, 1857 | S | O |
| **BOTHIDAE** |  |  |
| *Bothus mancus* Broussonet, 1782 | S | DC |
| *Bothus pantherinus* Rüppell, 1830 | S | NC |
| **CAESIONIDAE** |  |  |
| *Caesio varilineata* Carpenter, 1987 | S | DPL |
| *Caesio xanthalytos* Holleman, Connell & Carpenter, 2013 | S | DPL* |
| *Caesio xanthonata* Bleeker, 1853 | S | DPL |
| *Pterocaesio marri* Schultz, Herald, Lachner, Welander & Woods, 1953 | S | DPL |
| *Pterocaesio tile* Cuvier & Valenciennes, 1830 | S | DPL |
| **CALLIONMYIDAE** |  |  |
| *Neosynchiropus stellatus* Smith, 1963 | S | DC |
| **CARANGIDAE** |  |  |
| *Alectis ciliaris* Bloch, 1787 | P | DC |
| *Alectis indica* Rüppell, 1830 | P | DC |
| *Caranx bucculentus* Alleyne & Macleay, 1877 | S | DC |
| *Caranx heberi* Bennett, 1830 | S | DC |
| *Caranx ignobilis* Forsskål, 1775 | S | DC |
| *Caranx melampygus* Cuvier, 1833 | S | DC |
| *Caranx sexfasciatus* Quoy & Gaimard, 1825 | S | Pi |
| *Elagatis bipinnulata* Quoy & Gaimard, 1825 | S | DC |
| *Gnathanodon speciosus* Forsskål, 1775 | S | DC |
| *Seriola lalandi* Valenciennes, 1833 | S | DC |
| **CARCHARHINIDAE** |  |  |
| *Carcharhinus amblyrhynchos* Bleeker, 1856 | S | Pi |
| *Carcharhinus leucas* Müller & Henle, 1839 | P | DC |
| *Carcharhinus limbatus* Müller & Henle, 1839 | S | Pi |
| *Carcharhinus melanopterus* Quoy & Gaimard, 1824 | S | Pi |
| *Carcharhinus obscurus* Lesueur, 1818 | S | DC |
| *Triaenodon obesus* Rüppell, 1837 | S | DC |
| **CENTRISCIDAE** |  |  |
| *Aeoliscus strigatus* Günther, 1861 | P | DC |
| **CHAETODONTIDAE** |  |  |
| *Chaetodon auriga* Forsskål, 1775 | S | BSI |
| *Chaetodon blackburnii* Desjardins, 1836 | S | BSI |
| *Chaetodon dolosus* Ahl, 1923 | S | BSI |
| *Chaetodon guttatissimus* Bennett, 1833 | S | BSI |
| *Chaetodon interruptus* Ahl, 1923 | S | BSI |
| *Chaetodon kleinii* Bloch, 1790 | S | BSI |
| *Chaetodon lineolatus* Cuvier, 1831 | S | BSI |
| *Chaetodon lunula* Lacepède, 1802 | S | BSI |
| *Chaetodon madagaskariensis* Ahl, 1923 | S | BSI |
| *Chaetodon melannotus* Bloch & Schneider, 1801 | S | BSI |
| *Chaetodon meyeri* Bloch & Schneider, 1801 | S | BSI |
| *Chaetodon trifascialis* Quoy & Gaimard, 1825 | S | BSI |
| *Chaetodon xanthurus* Bleeker, 1857 | S | BSI |
| *Forcipiger flavissimus* Jordan & McGregor, 1898 | S | BSI |
| *Hemitaurichthys zoster* Bennett, 1831 | S | DPL |
| *Heniochus acuminatus* Linnaeus, 1758 | S | BSI |
| *Heniochus diphreutes* Jordan, 1903 | S | DPL |
| *Heniochus monoceros* Cuvier, 1831 | S | BSI |
| **CIRRHITIDAE** |  |  |
| *Cirrhitichthys oxycephalus* Bleeker, 1855 | S | DC |
| *Cyprinocirrhites polyactis* Bleeker, 1874 | S | DPL |
| *Oxycirrhites typus* Bleeker, 1857 | P | DPL |
| *Paracirrhites arcatus* Cuvier, 1829 | S | DC |
| *Paracirrhites forsteri* Schneider, 1801 | S | DC |
| CLINIDAE |  |  |
| *Clinus venustris* Gilchrist & Thompson, 1908 | S | NA |
| *Pavoclinus laurentii* Gilchrist & Thompson, 1908 | S | NA |
| CLUPEIDAE |  |  |
| *Gilchristella aestuaria* Gilchrist, 1913 | S | DPL |
| CONGRIDAE |  |  |
| *Heteroconger hassi* Klausewitz & Eibl-Eibesfeldt, 1959 | S | NC |
| DACTYLOPTERIDAE |  |  |
| *Dactyloptena orientalis* Cuvier, 1829 | S | NC |
| DASYATIDAE |  |  |
| *Dasyatis microps* Annandale, 1908 | S | NC* |
| *Himantura jenkinsii* Annandale, 1909 | S | NC |
| *Himantura uarnak* Gmelin, 1789 | S | NC |
| *Neotrygon kuhlii* Müller & Henle, 1841 | S | NC |
| *Taeniura lymma* Forsskål, 1775 | P | NC |
| *Taeniura meyeni* Müller & Henle, 1841 | S | NC |
| DIODONTIDAE |  |  |
| *Diodon holocanthus* Linnaeus, 1758 | S | NC |
| *Diodon hystrix* Linnaeus, 1758 | S | NC |
| *Diodon liturosus* Shaw, 1804 | S | NC |
| ECHENEIDAE |  |  |
| *Echeneis naucrates* Linnaeus, 1758 | S | NC |
| ENGRAULIDAE |  |  |
| *Thryssa vitrirostris* Gilchrist & Thompson, 1908 | S | DPL |
| EPHIPPIDAE |  |  |
| *Platax teira* Forsskål, 1775 | S | O |
| FISTULARIIDAE |  |  |
| *Fistularia commersonii* Rüppell, 1838 | S | Pi |
| GERREIDAE |  |  |
| *Gerres longirostris* Lacepède, 1801 | S | DC |
| GINGLYMOSTOMATIDAE |  |  |
| *Nebrius ferrugineus* Lesson, 1831 | P | NC |

| **FAMILIES - Species - Authors** | **Sighting**<br>**Record** | **Trophic**<br>**Category** |
| --- | --- | --- |
| **GOBIIDAE** |  |  |
| *Amblyeleotris steinitzi* Klausewitz, 1974 | S | DC |
| *Amblyeleotris wheeleri* Polunin & Lubbock, 1977 | S | DC* |
| *Caffrogobius saldanha* Barnard, 1927 | S | NA |
| *Valenciennea strigata* Broussonet, 1782 | S | DC |
| **HAEMULIDAE** |  |  |
| *Diagramma pictum* Thunberg, 1792 | S | DC |
| *Plectorhinchus flavomaculatus* Cuvier, 1830 | S | NC |
| *Plectorhinchus gaterinus* Forsskål, 1775 | S | NC |
| *Plectorhinchus playfairi* Pellegrin, 1914 | S | DC |
| *Plectorhinchus vittatus* Linnaeus, 1758 | S | NC |
| **HEMIRAMPHIDAE** |  |  |
| *Hyporhamphus affinis* Günther, 1866 | S | O |
| **HOLOCENTRIDAE** |  |  |
| *Myripristis adusta* Bleeker, 1853 | S | NPL |
| *Myripristis berndti* Jordan & Evermann, 1903 | S | NC |
| *Myripristis botche* Cuvier, 1829 | S | NC |
| *Myripristis murdjan* Forsskål, 1775 | S | NPL |
| *Myripristis vittata* Valenciennes, 1831 | S | NPL |
| *Neoniphon samara* Forsskål, 1775 | S | NC |
| *Pagellus natalensis* Steindachner, 1903 | S | O |
| *Sargocentron caudimaculatum* Rüppell, 1838 | S | NC |
| *Sargocentron diadema* Lacepède, 1802 | S | NC |
| *Sargocentron spiniferum* Forsskål, 1775 | S | NC |
| **ISTIOPHORIDAE** |  |  |
| *Istiompax indica* Cuvier, 1832 | S | Pi |
| *Istiophorus platypterus* Shaw, 1792 | P | Pi |
| *Makaira nigricans* Lacepède, 1802 | P | Pi |
| **KYPHOSIDAE** |  |  |
| *Kyphosus vaigiensis* Quoy & Gaimard, 1825 | S | H |
| **LABRIDAE** |  |  |
| *Anampses meleagrides* Valenciennes, 1840 | S | DC |
| *Bodianus anthioides* Bennett, 1832 | S | DC |
| *Bodianus axillaris* Bennett, 1832 | S | DC |
| *Bodianus diana* Lacepède, 1801 | S | DC |

| **FAMILIES - Species - Authors** | **Sighting Record** | **Trophic Category** |
| --- | --- | --- |
| *Bodianus trilineatus* Fowler, 1934 | S | DC* |
| *Anampses twistii* Bleeker, 1856 | S | DC |
| *Cheilinus trilobatus* Lacepède, 1801 | S | DC |
| *Cheilinus undulates* Rüppell, 1835 | S | DC |
| *Cheilio inermis* Forsskål, 1775 | S | DC |
| *Coris aygula* Lacepède, 1801 | S | DC |
| *Coris caudimacula* Quoy & Gaimard, 1834 | S | DC |
| *Coris cuvieri* Bennett, 1831 | S | DC |
| *Coris formosa* Bennett, 1830 | S | DC |
| *Gomphosus caeruleus* Lacepède, 1801 | S | DC |
| *Gomphosus varius* Lacepède, 1801 | S | DC |
| *Halichoeres cosmetus* Randall & Smith, 1982 | S | DC |
| *Halichoeres hortulanus* Lacepède, 1801 | S | DC |
| *Halichoeres iridis* Randall & Smith, 1982 | S | DC |
| *Halichoeres lapillus* Smith, 1947 | S | DC |
| *Halichoeres nebulosus* Valenciennes, 1839 | S | DC |
| *Halichoeres scapularis* Bennett, 1832 | S | DC |
| *Halichoeres zeylonicus* Bennett, 1833 | S | DC |
| *Halichoeres zulu* Randall & King, 2010 | S | DC |
| *Labroides bicolor* Fowler & Bean, 1928 | S | DC |
| *Labroides dimidiatus* Valenciennes, 1839 | S | DC |
| *Macropharyngodon bipartitus* Smith, 1957 | S | DC |
| *Macropharyngodon cyanoguttatus* Randall, 1978 | S | DC* |
| *Novaculichthys taeniourus* Lacepède, 1801 | S | DC |
| *Pseudocoris heteroptera* Bleeker, 1857 | S | DC |
| *Thalassoma amblycephalum* Bleeker, 1856 | S | DC |
| *Thalassoma hebraicum* Lacepède, 1801 | S | DC |
| *Thalassoma lunare* Linnaeus, 1758 | S | DC |
| **LUTJANIDAE** |  |  |
| *Aprion virescens* Valenciennes, 1830 | S | Pi |
| *Lutjanus ehrenbergii* Peters, 1869 | S | NC |
| *Lutjanus fulviflamma* Forsskål, 1775 | S | NC |
| *Lutjanus gibbus* Forsskål, 1775 | S | NC |
| *Lutjanus kasmira* Forsskål, 1775 | S | NC |
| *Lutjanus lutjanus* Bloch, 1790 | S | NC |
| *Lutjanus monostigma* Cuvier, 1828 | S | NC |
| *Lutjanus notatus* Cuvier, 1828 | S | NC |
| *Lutjanus rivulatus* Cuvier, 1828 | S | NC |
| *Lutjanus sebae* Cuvier, 1816 | S | NC |
| *Macolor niger* Forsskål, 1775 | S | NC |
| *Paracaesio sordida* Abe & Shinohara, 1962 | S | DPL |
| MALACANTHIDAE |  |  |
| *Malacanthus brevirostris* Guichenot, 1848 | S | DC |
| MICRODESMIDAE |  |  |
| *Nemateleotris magnifica* Fowler, 1938 | S | NPL |
| *Ptereleotris evides* Jordan & Hubbs, 1925 | S | NPL |
| *Ptereleotris heteroptera* Bleeker, 1855 | S | DPL |
| MOLIDAE |  |  |
| *Mola mola* Linnaeus, 1758 | P | DC |
| MONACANTHIDAE |  |  |
| *Aluterus scriptus* Osbeck, 1765 | S | O |
| *Acreichthys tomentosus* Linnaeus, 1758 | S | DC |
| *Cantherhines fronticinctus* Günther, 1867 | S | BSI |
| *Cantherhines pardalis* Rüppell, 1837 | S | BSI |
| *Pervagor janthinosoma* Bleeker, 1854 | S | NA |
| *Stephanolepis auratus* Castelnau, 1861 | S | NA |
| MONOCENTRIDAE |  |  |
| *Cleidopus gloriamaris* De Vis, 1882 | P | NA |
| MONODACTYLIDAE |  |  |
| *Monodactylus argenteus* Linnaeus, 1758 | S | DPL |
| MULLIDAE |  |  |
| *Mulloidichthys ayliffe* Uiblein, 2011 | S | NC |
| *Mulloidichthys flavolineatus* Lacepède, 1801 | S | NC |
| *Mulloidichthys vanicolensis* Valenciennes, 1831 | S | NC |
| *Parupeneus barberinus* Lacepède, 1801 | S | DC |
| *Parupeneus indicus* Shaw, 1803 | S | DC |
| *Parupeneus macronemus* Lacepède, 1801 | S | DC |
| *Parupeneus trifasciatus* Lacepède, 1801 | S | DC |
| MURAENIDAE |  |  |
| *Echidna nebulosa* Ahl, 1789 | S | NC |
| *Enchelycore pardalis* Temminck & Schlegel, 1846 | S | Pi |
| *Gymnomuraena zebra* Shaw, 1797 | S | NC |
| *Gymnothorax breedeni* McCosker & Randall, 1977 | S | NC |
| *Gymnothorax eurostus* Abbott, 1860 | S | NC |
| *Gymnothorax favagineus* Bloch & Schneider, 1801 | S | NC |
| *Gymnothorax flavimarginatus* Rüppell, 1830 | S | Pi |
| *Gymnothorax griseus* Lacepède, 1803 | S | NC* |
| *Gymnothorax javanicus* Bleeker, 1859 | S | NC |
| *Gymnothorax meleagris* Shaw, 1795 | S | DC |
| *Gymnothorax miliaris* Kaup, 1856 | S | DC |
| *Gymnothorax nudivomer* Günther, 1867 | S | NC* |
| *Gymnothorax undulates* Lacepède, 1803 | S | NC |
| *Rhinomuraena quaesita* Garman, 1888 | P | Pi |
| **MYLIOBATIDAE** |  |  |
| *Aetobatus narinari* Euphrasen, 1790 | P | DC |
| *Manta alfredi* Krefft, 1868 | S | DPL |
| *Manta birostris* Walbaum, 1792 | S | DPL |
| *Mobula japonica* Müller & Henle, 1841 | S | DPL |
| **ODONTASIPSIDAE** |  |  |
| *Carcharias taurus* Rafinesque, 1810 | S | DC |
| **OPHICHTHIDAE** |  |  |
| *Myrichthys colubrinus* Boddaert, 1781 | S | NC |
| *Myrichthys maculosus* Cuvier, 1816 | S | NC |
| *Pisodonophis cancrivorus* Richardson, 1848 | P | NC |
| **OPLEGNATHIDAE** |  |  |
| *Oplegnathus robinsoni* Regan, 1916 | S | O |
| **OSTRACIIDAE** |  |  |
| *Lactoria fornasini* Bianconi, 1846 | S | BSI* |
| *Lactoria cornuta* Linnaeus, 1758 | S | BSI |
| *Ostracion cubicus* Linnaeus, 1758 | S | BSI |
| *Ostracion meleagris* Shaw, 1796 | S | BSI |
| **PEGASIDAE** |  |  |
| *Eurypegasus draconis* Linnaeus, 1766 | S | BSI |
| **PEMPHERIDAE** |  |  |
| *Parapriacanthus ransonneti* Steindachner, 1870 | S | NPL |
| *Pempheris schwenkii* Bleeker, 1855 | S | NPL |
| PINGUIPEDIDAE |  |  |
| *Parapercis schauinslandii* Steindachner, 1900 | S | DC |
| PLATYCEPHALIDAE |  |  |
| *Papilloculiceps longiceps* Cuvier, 1829 | S | DC |
| PLOTOSIDAE |  |  |
| *Plotosus lineatus* Thunberg, 1787 | S | NC |
| POMACANTHIDAE |  |  |
| *Apolemichthys trimaculatus* Cuvier, 1831 | S | O |
| *Centropyge acanthops* Norman, 1922 | S | O |
| *Centropyge bispinosa* Günther, 1860 | S | O |
| *Centropyge multispinis* Playfair, 1867 | S | O |
| *Pomacanthus chrysurus* Cuvier, 1831 | S | O |
| *Pomacanthus imperator* Bloch, 1787 | S | O |
| *Pomacanthus rhomboides* Gilchrist & Thompson, 1908 | S | O* |
| *Pomacanthus semicirculatus* Cuvier, 1831 | S | BSI |
| *Pygoplites diacanthus* Boddaert, 1772 | S | BSI |
| POMACENTRIDAE |  |  |
| *Abudefduf natalensis* Hensley & Randall, 1983 | S | O |
| *Abudefduf sexfasciatus* Lacepède, 1801 | S | O |
| *Abudefduf vaigiensis* Quoy & Gaimard, 1825 | S | O |
| *Amphiprion allardi* Klausewitz, 1970 | S | O |
| *Amphiprion perideraion* Bleeker, 1855 | S | O* |
| *Chromis fieldi* Randall & DiBattista, 2013 | S | DPL |
| *Chromis nigrura* Smith, 1960 | S | DPL |
| *Chromis opercularis* Günther, 1867 | S | DPL |
| *Chromis viridis* Cuvier, 1830 | S | O |
| *Chromis weberi* Fowler & Bean, 1928 | S | DPL |
| *Chrysiptera brownriggii* Bennett, 1828 | S | O |
| *Chrysiptera unimaculata* Cuvier, 1830 | S | O |
| *Dascyllus aruanus* Linnaeus, 1758 | S | DPL |
| *Dascyllus carneus* Fischer, 1885 | S | O |
| *Dascyllus trimaculatus* Rüppell, 1829 | S | DPL |
| *Neopomacentrus cyanomos* Bleeker, 1856 | S | NA |
| *Plectroglyphidodon dickii* Liénard, 1839 | S | O |
| *Pomacentrus caeruleus* Quoy & Gaimard, 1825 | S | O |
| *Pomacentrus pavo* Bloch, 1787 | S | O |
| *Stegastes fasciolatus* Ogilby, 1889 | S | H |
| *Stegastes pelicier* Allen & Emery, 1985 | S | H |
| **PRIACANTHIDAE** |  |  |
| *Priacanthus hamrur* Forsskål, 1775 | S | NC |
| **PSEUDOCROMIDAE** |  |  |
| *Pseudochromis dutoiti* Smith, 1955 | S | DC |
| **RACHYCENTRIDAE** |  |  |
| *Rachycentron canadum* Linnaeus, 1766 | S | DC |
| **RHINCODONTIDAE** |  |  |
| *Rhincodon typus* Smith, 1828 | S | DPL |
| **RHINIDAE** |  |  |
| *Rhina ancylostoma* Bloch & Schneider, 1801 | P | NC |
| **RHINOBATIDAE** |  |  |
| *Rhinobatus annulatus* Müller & Henle, 1841 | P | NC |
| *Rhinobatus leucospilus* Norman, 1926 | S | NC |
| *Rhynchobatus djiddensis* Forsskål, 1775 | S | NC |
| **SCARIDAE** |  |  |
| *Chlorurus cyanescens* Valenciennes, 1840 | S | H |
| *Chlorurus sordidus* Forsskål, 1775 | S | H |
| *Scarus ghobban* Forsskål, 1775 | S | H |
| *Scarus rubroviolaceus* Bleeker, 1847 | S | H |
| *Scarus scaber* Valenciennes, 1840 | S | H |
| *Scarus tricolor* Bleeker, 1847 | S | H |
| **SCOMBRIDAE** |  |  |
| *Euthynnus affinis* Cantor, 1849 | S | DC |
| *Gymnosarda unicolor* Rüppell, 1836 | S | Pi |
| *Katsuwonus pelamis* Linnaeus, 1758 | S | DC |
| *Scomberomorus commerson* Lacepède, 1801 | S | Pi |
| *Scomberomorus plurilineatus* Fourmanoir, 1966 | P | Pi |
| *Thunnus albacares* Bonnaterre, 1788 | S | DC |
| **SCORPAENIDAE** |  |  |
| *Caracanthus maculatus* Gray, 1831 | S | NA |
| *Dendrochirus brachypterus* Cuvier, 1829 | S | NC |
| *Dendrochirus zebra* Cuvier, 1829 | S | NC |
| *Parascorpaena mossambica* Peters, 1855 | S | NA |
| *Pterois antennata* Bloch, 1787 | S | DC |
| *Pterois miles* Bennett, 1828 | S | Pi |
| *Rhinopias eschmeyeri* Condé, 1977 | P | Pi* |
| *Rhinopias frondosa* Günther, 1892 | P | Pi |
| *Scorpaenopsis diabolus* Cuvier, 1829 | S | Pi |
| *Scorpaenopsis oxycephala* Bleeker, 1849 | S | Pi |
| *Scorpaenopsis venosa* Cuvier, 1829 | S | DC |
| *Sebastapistes cyanostigma* Bleeker, 1856 | S | NA |
| *Taenianotus triacanthus* Lacepède, 1802 | S | DC |
| **SERRANIDAE** |  |  |
| *Cephalopholis argus* Schneider, 1801 | S | Pi |
| *Cephalopholis miniata* Forsskål, 1775 | S | NC |
| *Cephalopholis sonnerati* Valenciennes, 1828 | S | NC |
| *Epinephelus chlorostigma* Valenciennes, 1828 | S | NC |
| *Epinephelus fasciatus* Forsskål, 1775 | S | NC |
| *Epinephelus flavocaeruleus* Lacepède, 1802 | P | Pi |
| *Epinephelus lanceolatus* Bloch, 1790 | P | NC |
| *Epinephelus macrospilos* Bleeker, 1855 | S | DC |
| *Epinephelus malabaricus* Bloch & Schneider, 1801 | S | NC |
| *Epinephelus merra* Bloch, 1793 | S | Pi |
| *Epinephelus rivulatus* Valenciennes, 1830 | S | Pi |
| *Epinephelus tauvina* Forsskål, 1775 | S | Pi |
| *Epinephelus tukula* Morgans, 1959 | S | NC |
| *Grammistes sexlineatus* Thunberg, 1792 | S | NC |
| *Nemanthias carberryi* Smith, 1954 | S | DPL |
| *Plectropomus punctatus* Quoy & Gaimard, 1824 | S | Pi |
| *Pogonoperca punctata* Valenciennes, 1830 | S | NC* |
| *Pseudanthias evansi* Smith, 1954 | S | DPL |
| *Pseudanthias squamipinnus* Peters, 1855 | S | DPL |
| **SIGANIDAE** |  |  |
| *Siganus luridus* Rüppell, 1829 | S | H |
| *Siganus sutor* Valenciennes, 1835 | S | H |
| **SOLEIDAE** |  |  |
| *Solea turbynei* Gilchrist, 1904 | S | NA |
| SPARIDAE |  |  |
| *Chrysoblephus puniceus* Gilchrist & Thompson, 1908 | S | DC |
| *Diplodus hottentotus* Smith, 1844 | S | DC |
| SPHRYNIDAE |  |  |
| *Sphyrna lewini* Griffith & Smith, 1834 | S | DC |
| SPHYRAENIDAE |  |  |
| *Sphyraena putnamae* Jordan & Seale, 1905 | S | NC |
| STEGOSTOMATIDAE |  |  |
| *Stegostoma fasciatum* Hermann, 1783 | S | NC |
| SYNANCEIIDAE |  |  |
| *Synanceia verrucosa* Bloch & Schneider, 1801 | S | Pi |
| SYNGNATHIDAE |  |  |
| *Corythoichthys intestinalis* Ramsay, 1881 | P | DC |
| *Doryrhamphus dactyliophorus* Bleeker, 1853 | S | DPL |
| *Hippocampus borboniensis* Duméril, 1870 | S | DPL* |
| *Hippocampus camelopardalis* Bianconi, 1854 | P | DPL* |
| *Hippocampus histrix* Kaup, 1856 | S | DPL |
| *Hippocampus kuda* Bleeker, 1852 | S | DPL |
| *Solenostomus cyanopterus* Bleeker, 1854 | S | DC |
| *Trachyrhamphus bicoarctatus* Bleeker, 1857 | S | NA |
| SYNODONTIDAE |  |  |
| *Synodus dermatogenys* Fowler, 1912 | S | Pi |
| *Synodus jaculum* Russell & Cressey, 1979 | S | Pi |
| TETRAODONTIDAE |  |  |
| *Arothron hispidus* Linnaeus, 1758 | S | NC |
| *Arothron meleagris* Anonymous, 1798 | S | NC |
| *Arothron nigropunctatus* Bloch & Schneider, 1801 | S | NC |
| *Arothron stellatus* Anonymous, 1798 | S | NC |
| *Canthigaster amboinensis* Bleeker, 1864 | S | H |
| *Canthigaster bennetti* Bleeker, 1854 | S | O |
| *Canthigaster janthinoptera* Bleeker, 1855 | S | O |
| *Canthigaster smithae* Allen & Randall, 1977 | S | O* |
| *Canthigaster solandri* Richardson, 1845 | S | O |
| *Canthigaster valentine* Bleeker, 1853 | S | O |
| TETRAROGIDAE |  |  |
| *Ablabys binotatus* Peters, 1855 | S | NA |
| *Ablabys macracanthus* Bleeker, 1852 | S | NA |
| TORPEDINIDAE |  |  |
| *Torpedo marmorata* Risso, 1810 | S | Pi |
| *Torpedo spp.* | S | Pi |
| ZANCLIDAE |  |  |
| *Zanclus cornutus* Linnaeus, 1758 | S | DC |
| **Trophic Categories: Herbivore (H); Omnivore (O); Browser of Sessile Invertebrates (BSI); Diurnal Carnivore (DC); Nocturnal Carnivore (NC); Piscivore (Pi); Diurnal Planktivore (DPL); Nocturnal Planktivore (NPL); Unknown (NA)** |  |  |

**Table 3.** The diversity of reef fish species and families from other areas in the southwestern Indian Ocean. SRobs = recorded species richness; SR_theor_ = theoretical species richness predicted by the Coral Fish Diversity Index (Allen & Werner, 2002).

| Location | Geographical Coordinates | SR<sub>obs</sub> | SR<sub>theor</sub> | No. of families | SR<sub>obs</sub> to no. of families ratio (2 d. p.) | Source |
| --- | --- | --- | --- | --- | --- | --- |
| Praia do Tofo & Praia da Barra | 23°51'S, 33°54'E | 353 | 329 | 79 | 4.47:1 | Present study |
| Bazaruto Archipelago National Park | 21°43'S, 35°27'E | 249 | 359 | 40 | 6.23:1 | Maggs *et al.* 2010 |
| Maputo Bay | 26°S, 32°54'E | 327 | 349 | 58 | 5.64:1 | Schleyer & Pereira, 2014 |
| Juan de Nova | 17°03'S, 42°43'E | 299 | 423 | 55 | 5.44:1 | Chabanet & Durville, 2005 |
| Andavadoaka | 22°05'S, 43°12'E | 334 | 430 | 58 | 5.76:1 | Gillibrand, Harries & Mara, 2007 |
| Glorieuses Islands | 11°33'S, 47°20'E | 332 | 451 | 57 | 5.82:1 | Durville, Chabanet & Quod, 2003 |
| St. Lucia Marine Reserve | 27°44'S, 32°40'E | 258 | 349 | 48 | 5.38:1 | Floros *et al.* 2012 |
| Mafia Island | 7°52'S, 39°45'E | 394 | 515 | 56 | 7.04:1 | Garpe & Ohman, 2003 |
| Europa Island | 22°21'S, 40°21'E | 389 | 468 | 62 | 6.27:1 | Fricke *et al.* 2013 |
| Ponta do Ouro Partial Marine Reserve | 26°27'S, 32°56'E | 376 | 318 | 90 | 4.18:1 | Pereira, Videira & Abrantes, 2004 |

## Discussion

This is the first assessment of ichthyofaunal diversity of the seas around Praia do Tofo and Praia da Barra in southern Mozambique. Through the use of underwater observations supplemented by past records, 353 species were recorded from the coral reefs spanning 40 km of the southern coastline of the Inhambane province. These results provide a higher estimation of fish species richness than is predicted by the Coral Fish Diversity Index. The diversity of the PTPB area is similar to that recorded in other areas of the southwestern Indian Ocean where visual observations have been the primary data collection method (Table 3) (Maggs *et al*., 2010; Chabanet & Durville, 2005; Gillibrand, Harries & Mara, 2007; Durville, Chabanet & Quod, 2003). In particular, SR_theor_ shows high similarity to areas in southern Mozambique and South Africa that are fully or partially protected (e.g. Floros *et al*. 2012; Maggs *et al*. 2010; Pereira, Videira & Abrantes, 2004).

The sub-tropical reefs of the PTPB area have levels of coral cover (Motta *et al*. 2002), which may be assumed to result in a low diversity of fish communities (Komyakova, Munday & Jones, 2013). However, the current study finds a relatively high ichthyofaunal species richness which is comparable to areas with higher coral cover (e.g. Gillibrand, Harries & Mara, 2007; Table 3). This may be partly explained by the extensive visual sampling design used. The high sampling time employed in this study (over 36 hours of underwater observations) allowed for the observation of some cryptic species that would be missed by shorter visual surveying. For example, four species of gobies and eight species of blennies were recorded on reefs of PTPB (Table 2). Therefore while visual censuses generally do not accurately capture the diversity of cryptobenthic species (Ackerman & Bellwood, 2000) this limitation can be reduced. A high number of families were also recorded in comparison to other areas in the region (Table 3), suggesting a high proportion of uncommon species were observed. The impact of greater sampling effort on species records is evident in the results of Gillibrand, Harries & Mara (2007). These authors examined a smaller area than the current study and recorded 334 species by conducting visual observations across a twelve month period. In contrast, Chabanet and Durville (2005) recorded more than 50 fewer species around Juan de Nova island through 30 hours of visual surveying. This highlights that sampling effort does not solely account for the high fish diversity recorded in the PTPB area.

The present study necessarily examined a large depth range (1-32 m) in order to capture the range of habitats present in the area. As such a higher number of specialist species are expected to have been identified due to the wider variety of physical habitats and biological conditions (Bridge *et al*. 2016; Jankowski, Graham & Jones, 2015), Significant changes in fish assemblages with depth have been observed in previous studies (e.g. Friedlander & Parrish, 1998) and this is likely to be the same in the current study. This may also explain the high number of families observed (Table 3).

Coastal upwelling in these seas drives high levels of primary productivity and in turn supports abundant populations of large charismatic species (Rohner *et al*. 2014). It is also likely to influence the reef fish diversity of the area, potentially boosting species richness in two ways. Firstly, cooler waters allow the area to support species more common in temperate waters (e.g. *Seriola lalandi*, *Oplegnathus robinsoni*). Anderson *et al*. (2015) proposed the appearance of species characteristic of higher latitudes in their sub-tropical study site to regions of cool water upwelling. In the current study water temperatures were recorded between 18-29°C and the influx of cool water may also influence diversity in the sub-tropical PTPB area. Secondly, upwelling supports high plankton abundance which can reduce competitive exclusion in planktivorous species (Abrams, 1995). This would allow the co-existence of more species on lower trophic levels, an effect which may then propagate up the food chain to produce a higher diversity of secondary and tertiary consumers. The relationship between primary productivity and diversity has been previously acknowledged (Waide *et al*. 1999).

This study demonstrates the PTPB area’s biological value beyond its resident megafauna populations, and the future for a broader value of ecotourism to the region. Whilst the relatively large sampling extent precludes comprehensive comparisons with other studies in the southwestern Indian Ocean, the results show that the coral reef ecosystem of PTPB hosts a reef fish community comparable to more isolated or protected areas. As such the current study suggests that the reefs of PTPB are in good condition, despite the large associated human population. Targeted research is needed to examine the current health status of these reefs and to provide a baseline for monitoring impacts of future expansion of tourism and fishing activities in the region.

## Acknowledgements

I would like to thank Peri Peri dive centre and the Underwater Africa volunteer program for their support in undertaking both SCUBA and snorkel surveys. Sincere thanks to Dr Tracy Ainsworth, Dr William Leggat, Dr Hudson Pinheiro and an anonymous reviewer for comments on and improvements to the manuscript. Finally, thank you to all those friends and strangers who provided photographic evidence of rare species.

## References

Abrams PA. 1995. Monotonic or unimodal diversity-productivity gradients: what does competition theory predict? Ecology, 76: 2019–2027. DOI: 10.2307/1941677

Ackerman JL & Bellwood DR. 2000. Reef fish assemblages: a re-evaluation using enclosed rotenone stations. Marine Ecology Progression Series, 206: 227–237.

Allen GR & Werner TB. 2002. Coral reef fish assessment in the ‘coral triangle’ of southeastern Asia. Environmental Biology of Fishes, 65: 209–214. DOI: 10.1023/A:1020093012502

Anderson AB, Carvalho-Filho A, Morais RA, Nunes LT, Quimbayo JP & Floeter SR. 2015. Brazilian tropical fishes in their southern limit of distribution: checklist of Santa Catarina’s rocky ichthyofauna reef, remarks and new records. Check List, 11: art1688. DOI: 10.15560/11.4.1688

Bellwood DR & Hughes TP. 2001. Regional-scale assembly rules and biodiversity of coral reefs. Science, 292: 1532–1534.

Bridge TCL, Luiz OJ, Coleman RR, Kane CN & Kosaki RK. 2016. Ecological and morphological traits predict depth-generalist fishes on coral reefs. Proceedings of the Royal Society B, 283: 20152332. DOI: 10.1098/rspb.2015.2332

Chabanet P & Durville P. 2005. Reef fish inventory of Juan de Nova’s natural park (Western Indian Ocean). Western Indian Ocean Journal of Marine Science, 4: 145–162. DOI: 10.4314/wiojms.v4i2.28484

Chabanet P, Tessier E, Durville P, Mulochau T & René F. 2002. Fish communities of the Geyser and Zélée coral banks (Western Indian Ocean). Cybium, 26: 11–26.

Cleland EE. 2011. Biodiversity and ecosystem stability. Nature Education Knowledge, 3: pp. 14.

Durville P, Chabanet P & Quod JP. 2003. Visual census of the reef fishes in the natural reserve of the Glorieuses Islands (Western Indian Ocean). Western Indian Ocean Journal of Marine Science, 2: 95–104.

Floros C, Schleyer M, Maggs JQ & Celliers L. 2012. Baseline assessment of high-latitude coral reef fish communities in southern Africa. African Journal of Marine Science, 34: 55–69. DOI: 10.2989/1814232X.2012.673284

Friedlander AM & Parrish JD. 1998. Habitat characteristics affecting fish assemblages on a Hawaiian coral reef. Journal of Experimental Marine Biology and Ecology, 224: 1–30. DOI: 10.1016/S0022–0981(97)00164–0

Garpe KC & Öhman MC. 2003. Coral and fish distribution patterns in Mafia Island Park Marine, Tanzania: fish-habitat interactions. Hydrobiologia, 498: 191–211. DOI: 10.1023/A:1026217201408

Gillibrand CJ, Harries AR & Mara E. 2007. Inventory and Spatial Assemblage Study of Reef Fish in the Area Andavadoaka of, South-West Madagascar (Western Indian Ocean). Western Indian Ocean Journal of Marine Science, 6: 183–197. DOI: 10.14314/wiojms.v612.48239

Harmelin-Vivien ML. 1979. Ichtyofaune des récifs coralliens en France Outre-Mer. ICRI. Doc. Secrétariat d’Etat à l’Outre-Mer et Ministère de l’Aménagement du Territoire et de l’Environment. pp 136.

Hiatt WR & Strasberg DW. 1960. Ecological relationship of the fish fauna on coral reefs of the Marshall Islands. Ecological Monograph, 30: 65–127

Hobson ES. 1974. Feeding relationships of teleostean fish on coral reefs Kona in, Hawaii. Fish Bulletin, 72: 915–1031

Instituto Nacional de Estatística. 2007. Recenseamento Geral da População e Habitação, Indicadores Socio-Demográficos: Província da Inhambane. 3o Censo Geral da População e Habitação: pp. 5.

Jankowski MW, Graham NAJ & Jones GP. 2015. Depth gradients diversity in, distribution and habitat specialisation in coral reef fishes: implications for the depth-refuge hypothesis. Marine Ecology Progression Series, 540: 203–215. DOI: 10.3354/meps11523

Komyakova V, Munday PL & Jones GP. 2013. Relative Importance of Cover Coral, Habitat Complexity and Diversity in Determining the Structure of Reef Fish Communities. PLoS One, 8: e83178. DOI: 10.1371/journal.pone.0083178

Kulbicki M. 1988. Patterns in the trophic structure of fish populations across the SW lagoon of New Caledonia. Proceedings of the 6th International Coral Reef Symposium, Townsville, Australia (August 8–12), 2: 305–312.

Maggs JQ, Floros C, Pereira MAM. & Schleyer MH. 2010. Rapid Visual Assessment of Fish Communities on Selected Reefs in the Bazaruto Archipelago. Western Indian Ocean Journal of Marine Science, 9; 115–134.

Motta H, Pereira MAM, Gonçalves M, Ridgway T & Schleyer MH. 2002. Coral reef monitoring in Mozambique (2000). MICOA/CORDIO/ORI/WWF. Maputo, Mozambique Coral Reef Management Programme.

Mutimucuio M & Meyer D. 2011. Pro-poor employment and procurement: a tourism value chain analysis of peninsula Inhambane, Mozambique. In: van der Duim R, Meyer D, Saarinen J & Zellmer K (eds.). New alliances tourism for, conservation and development in Eastern and Southern Africa. Eburon, Delft.

Myers RF. 1999. Micronesian reef fishes. Guam: Coral Graphics. 298pp.

Pereira MAM. 2000. Preliminary checklist of reef-associated fishes of Mozambique. MICOAE, Maputo, pp. 21.

Pierce SJ, Méndez-Jiménez A, Collins K, Rosero-Caicedo M & Monadjem A. 2010. Developing a Code of Conduct for whale shark interactions in Mozambique. Aquatic Conservation: Marine and Freshwater Ecosystems, 20: 782–788. DOI: 10.1002/aqc.1149

Rohner CA, Pierce SJ, Marshall AD, Weeks SJ, Bennett MB & Richardson AJ. 2013. Trends in sightings and environmental influences on a coastal aggregation of manta rays and whale sharks. Marine Ecology Progression Series, 482: 153–168. DOI: 10.3354/meps10290

Rohner CA, Weeks SJ, Richardson AJ, Pierce SJ, Magno-Canto MM, Feldman GC, Cliff G & Roberts MJ. 2014. Oceanographic influences on a global whale shark hotspot in southern Mozambique. PeerJ PrePrints, 2:e661v1. DOI: 10.7287/peerj.preprints.661v1

Schleyer MH & Celliers L. 2005. The coral reefs of Island Bazaruto, Mozambique, with recommendations for their management. Western Indian Ocean Journal of Marine Science, 4: 227–236. DOI: 10.4314/wiojms.v4i2.28492

Tibiriçá Y, Birtles A, Valentine P & Miller DK. 2011. Diving Tourism in Mozambique: An Opportunity at Risk? Marine Environments, 7: 141–151. DOI: 10.3727/154427311X13195453162732

United Nations & World Heritage Convention. 2014. Assessing marine world heritage from an ecosystem perspective. The Western Indian Ocean, UN: 71–92 pp.

Van der Elst RP & Everett BI. 2015. Offshore fisheries of the Southwest Indian Ocean: their status and the impact on vulnerable species. Oceanographic Institute Research, Special Publication, 10: 448pp.

Venables S, Winstanley G, Bowles L & Marshall AD. 2016. A giant opportunity: the economic impact of manta rays on the Mozambican tourism industry – an incentive for increased management and protection. Tourism in Marine Environments¸ 12: 51–68. DOI: 10.3727/154427316X693225

Waide RB, Willig MR, Steiner CF, Mittelbach G, Gough L, Dodson SI, Juday GP & Parmenter R. 1999. The relationship between productivity and species richness. Annual Review of Ecology and Systematics, 30: 257–300.

Watson M, Righton D, Austin T & Ormond R. 1996. The effects of fishing on coral reef abundance and diversity. Journal of the Marine Biological Association of the United Kingdom, 76: 29–233. DOI: 10.1017/S0025315400029179

Wickel J, Jamon A, Pinault M, Durville P & Chabanet P. 2014. Species composition and structure of marine fish communities of Mayotte Island (south-western Indian Ocean). Cybium, 38: 179–203. DOI: 10.1016/j.biocon.2013.12.0290006–3207

